# Glial cell states bias the regeneration of neuron types across the newt life cycle

**DOI:** 10.64898/2026.05.26.728055

**Authors:** Alonso Ortega-Gurrola, Mayra Kalaora, Dreyton Amador, Jamie Woych, Maria Antonietta Tosches

**Author notes:** Correspondence should be addressed to M.A.T.

## Abstract

Salamanders have outstanding regenerative abilities, which tend to decline in post-metamorphic life stages. Among various tissues, these amphibians can regenerate the brain from ependymoglia cells, an adult neural stem cell population. Ependymoglia cells are heterogeneous; yet, whether ependymoglia cell diversity underlies variation of regenerative capacity across brain regions and life cycle stages remains poorly studied. Here we present a cell type comparison of regeneration in the pallium (dorsal telencephalon) of pre- and post-metamorphic newts. We found that ependymoglia cells exist in a continuum of cell states ranging from active proliferation to quiescence across life cycle stages, with a deep quiescence state featuring expression of mammalian astrocyte genes. Ependymoglia cell state changes are associated with a slower onset of proliferation and neurogenesis in post-metamorphic animals. Comparisons with developmental and adult neurogenesis reveal that pallial ependymoglia cells retain regional restrictions but can override temporal fate restrictions in response to an injury, producing neurons that are normally born only in early development. We thus find that brain regeneration in newts is not a simple amplification of adult neurogenesis, but a distinct process where the initial molecular state of ependymoglia cells biases the relative proportions of regenerated neuron types. Our findings establish post-metamorphic newts as a system to study how astrocyte-like glial cells can activate a neurogenic program in response to brain injury.

## Introduction

Vertebrates differ markedly in their capacity to regenerate the brain after an injury (Tanaka & Ferretti, 2009). In non-regenerative species like mammals, mechanical damage causes the formation of scar tissue and irreversible neuronal loss. Conversely, in highly regenerative fish and amphibian species, brain damage is followed by a neurogenic response and full repair (Lust & Tanaka, 2019). Yet, the variables that underlie these dramatic differences in regenerative capacity remain elusive.

In the brain and other regions of the central nervous system (CNS), regeneration is sustained by glial cells with neural stem cells properties that resume proliferation and enter neurogenesis in response to injury. In the zebrafish retina, Müller glia cells dedifferentiate and produce all the major retinal cell types (Lyu et al., 2023). In the fish and amphibian brain and spinal cord, ependymoglia cells (EGCs) are the main source of regenerated neurons (Berg et al., 2011; Fu et al., 2026; Goldshmit et al., 2012; Hui et al., 2010; Kaslin et al., 2013; Maden et al., 2013; Mchedlishvili et al., 2012; Parish et al., 2007; Rodrigo Albors et al., 2015). These cells are described as GFAP-expressing glial cells that retain contact to the ventricle and extend radial processes to the pial surface (Becker & Becker, 2015). EGCs exhibit properties of mammalian astrocytes (Chen et al., 2020; Morizet et al., 2024), neural stem cells (NSCs) (Lust et al., 2022; Morizet et al., 2024), and ependymal cells (Lust et al., 2022). Like their mammalian counterparts, EGCs originate at the end of developmental neurogenesis from radial glia cells (RGCs) that exit the cell cycle and activate the expression of glia differentiation genes (Deryckere et al., 2025; Joven et al., 2018). In the adult brain, EGCs occasionally reenter the cell cycle and produce new neurons, sustaining adult (homeostatic) neurogenesis. The widespread presence of EGCs is thus considered a key prerequisite for brain regeneration (Alunni & Bally-Cuif, 2016; Fu et al., 2026; Kirkham et al., 2014; Kroehne et al., 2011). However, the mere presence of EGCs is not sufficient to explain the variation of regenerative capacity in vertebrates: for example, frogs also have EGCs, but their brains do not regenerate after metamorphosis (Filoni et al., 1995; Yoshino & Tochinai, 2004). The relative inability of a species to regenerate despite having EGCs suggests that EGCs may exist in various cellular states or subtypes with distinct neurogenic properties. Accordingly, the distribution of these states or subtypes across species, or even life-cycle stages of the same species, may potentially explain variation of regenerative capacity.

Molecular studies have indeed provided some initial evidence of EGCs heterogeneity in anamniotes. Transcriptomic analyses in zebrafish revealed substantial heterogeneity and subsets of astrocyte-like EGCs (Cosacak et al., 2019; Lange et al., 2020; Morizet et al., 2024, 2025). In the killifish, glial transcriptomic heterogeneity is associated not only with distinct spatial locations but also with aging (Ayana et al., 2024). In the red spotted newt, telencephalic EGCs display region-specific proliferation dynamics (Kirkham et al., 2014). However, two key questions remain unanswered. First, it is unclear how EGCs types or states change over time, as animals progress from larval to adult stages. Second, it is largely unknown whether EGCs types or states respond differently to brain injury and thus influence regenerative capacity or fidelity. For example, EGCs may differ in their proliferation rate or pattern, or they may have different differentiation capacity. In the mammalian brain, adult neural stem cells are lineage-restricted and produce only a limited set of neuron types. Whether EGCs are also lineage-restricted in the brains of adult fish and amphibians, leading to incomplete or limited regeneration at the cell type level, remains largely untested.

In order to answer these questions, it is necessary to characterize EGC diversity and regenerative responses in a tractable system with clearly separated developmental and adult life stages. It is further crucial to define, at the cell type resolution, the repertoire of neurons generated by EGCs during adult neurogenesis and following injury, and determine whether EGCs have a limited neurogenic potential or can recapitulate developmental neurogenesis. Among amphibians, brain regeneration at cell type resolution has so far been studied primarily in the axolotl, a salamander species that retains larval characteristics throughout life and does not undergo metamorphosis (Amamoto et al., 2016; Lust et al., 2022; Wei et al., 2022). By contrast, newts have a complete life cycle including a post-metamorphic adult stage and also display robust regenerative abilities (Berg et al., 2011; Parish et al., 2007; Urata et al., 2018); however, brain regeneration has been poorly investigated with cell type resolution in newts and in other amphibians with a complete life cycle.

Here we analyze adult neurogenesis and regeneration in the Iberian ribbed newt *Pleurodeles waltl* (Joven et al., 2019; Matheson et al., 2025). With its distinct pre- and post-metamorphic stages, this species allows studying differences between regeneration and development. Building on previous work on the *Pleurodeles* pallium, the dorsal part of the telencephalon (Deryckere et al., 2025; Woych et al., 2022), we compare the transcriptomes of pallial neural progenitors across life cycle stages and characterize their responses to mechanical injury of the dorsal pallium (DP). Our analyses reveal that pallial EGCs exist in different quiescence states across stages and pallial regions, with deep quiescent EGCs characterized by the expression of mammalian astrocyte genes. In adults, the regenerative response is slower compared to larvae, in line with the presence of EGCs in deep quiescence. However, we find that regeneration, but not adult neurogenesis, produces all the neuron types originally present in the pallium, including those that are normally generated only during early development. Taken together, these findings demonstrate that adult neurogenesis and regeneration are two distinct processes and show that molecular heterogeneity among EGCs influences regenerative outcomes after brain injury in newts.

## Results

### Transcriptomic states of *Pleurodeles* neural stem cells across telencephalic regions and life cycle stages

In the newt pallium, neuron types are arranged along two orthogonal axes, established during development by spatial and temporal patterning of radial glia progenitors (Deryckere et al., 2025). Spatial patterning defines molecularly-distinct regions along the medio-lateral axis: the medial, dorsal, lateral, and ventral pallium, plus the pallial amygdala (MP, DP, LP, and VP, and AMY, respectively) (**Figure 1A**) (Deryckere et al., 2025; Woych et al., 2022). In some of these regions, including the DP, RGCs are multipotent and, over time, sequentially produce two distinct neuronal layers: superficial early-born neurons, generated before larval stage 50, and deep late-born neurons, born after stage 50 (Deryckere et al., 2025) (**Figure 1A**). At the end of the larval period, RGCs mature into EGCs (**Figure 1B**), and neurogenesis persists in the adult newt brain, although with regional-specific rates (Joven et al., 2018). We thus asked whether molecular differences of neural stem cells (RGCs iand EGCs) may underlie differences between larval and adult neurogenesis. To do so, we leveraged previous and new scRNA-seq data from the brains of larvae (eight developmental stages (Deryckere et al., 2025)) and adult animals (after expansion of the (Woych et al., 2022) dataset), which we re-aligned to the latest genome reference (Brown et al., 2025) (**Supplementary Figure 1A-B**). After quality filtering, we obtained 67,541 cells in the adult dataset and 132,548 cells in the developmental dataset (**Figure 1C**). We then annotated both datasets by performing label transfer from our previous transcriptomic-aligned annotations (**Supplementary Figure 1A-B**, see Methods).

**Figure 1.**
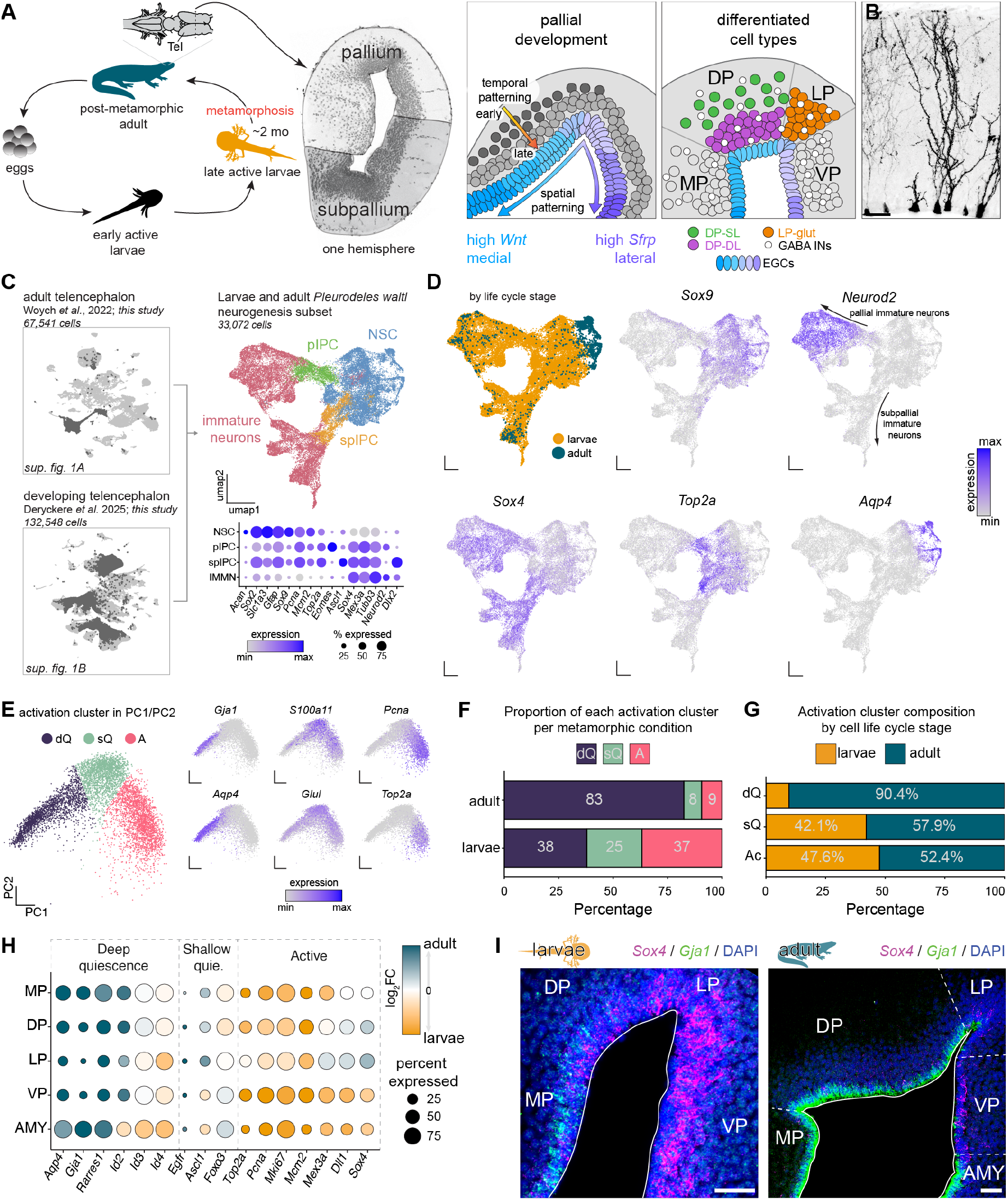
Newt EGCs occupy distinct transcriptomic states with region-specific distributions across the pallium. (A) Left: Life cycle of the Iberian ribbed newt *Pleurodeles waltl*. Right: schematic summarizing how the temporal and spatial patterning of radial glia progenitors in the developing pallium produce neuronal diversity along the medio-lateral and radial axes. (B) Morphology of newt ependymoglia cells labeled by electroporation of a CAG::GFP plasmid in the adult. (C) Left: UMAP plots of scRNAseq datasets from the adult (top) and developing (bottom) newt brain, highlighting the telencephalic cells that were used for subsequent analysis (see also Figure S1). Top right: UMAP plot of the neurogenic trajectory, showing neural progenitors and immature neurons pooled across larval and adult datasets. Top bottom: expression of genes defining major cell classes in the neurogenic trajectory: neural stem cells (NSCs), which include both RGCs and EGCs; pallial and subpallial intermediate progenitor cells (pIPCs and spIPCs, respectively), and immature neurons. Dot size indicates percentage of cells in the cluster expressing the gene, dot color indicates expression level. (D) UMAP plots of the neurogenic trajectory, colored according to life cycle stage, and to the expression of key marker genes in the trajectory. (E) Left: PCA plot of pallial progenitors (RGCs and EGCs) colored based on discrete assignments to cell state clusters (see also Figure S2). Right: expression of genes indicating active proliferation or quiescence in PCA space. (F) Stacked barplot showing the proportions of progenitors within each cell state cluster among adult EGCs and stage 55 larvae RGCs. (G) Stacked barplot showing the composition of the cell state clusters according by life cycle stage. (H) Expression of genes associated to quiescence and activation of NSCs by pallial region. Dot size indicates percentage of cells in the pallial region expressing the gene, dot color indicates log2 fold change of expression between adult EGCs and stage 55 larval cells. See Figure S1 for the identification of pallial regions. (I) Coronal sections at mid-telencephalic level showing the expression of the deep quiescence EGC marker *Gja1* and the immature neurons marker *Sox4* in stage 55 larvae (left) and in adults (right). Scale bar: 50 µm. Abbreviations: DP-DL, dorsal pallium deep-layer neurons; DP, dorsal pallium; EGC, ependymoglia cells; GABA, GABAergic; glut, glutamatergic; INs, interneurons; pIPC, pallial intermediate progenitor cell; spIPC, subpallial intermediate progenitor cell; AMY, amygdala; LP, lateral pallium; mo, months old; MP, medial pallium; NSC, neural stem cell; DP-SL, dorsal pallium superficial-layer neurons; Tel, telencephalon; VP, ventral pallium.

Once both adult and larval datasets were annotated, we selected and then merged telencephalic progenitors and immature neurons. Telencephalic cells were identified as *Foxg1* expressing cells; RGCs, EGCs and intermediate progenitor cells (IPCs) (Deryckere et al., 2025) from the high expression of *Sox9, Slc1a3, Gfap, Sox2*, and *Fabp7*; immature neurons from the high expression of *Sox4* and *Mex3a* and low expression of differentiation markers such as *Snap25* and *Syt1*. The final dataset contained 33,072 cells, capturing the neurogenic trajectories from RGCs and EGCs to pallial and subpallial immature neurons (**Figure 1C**). On the basis of gene expression and cell cycle phase (Deryckere et al., 2025), we identified pallial (*Eomes*+) and subpallial (*Ascl1*+) IPCs in both the developmental stages as well as in the adult (**Figure 1C-D**). Interestingly, neural stem cells were molecularly heterogeneous, with a major split between cells expressing high levels of cell cycle genes like *Top2a*, and cells expressing mature glia genes such as *Aqp4* and *Acan* (**Figure 1C-D**). This split correlated with the origin of cells from larval and adult brains, respectively. Finally, to further characterize pallial neurogenesis, we selected pallial progenitors defined as *Emx1* and *Pax6* expressing-cells (see Methods).

We then further explored the pallial RGCs/EGCs dataset (6,959 cells, of which 2,214 were from adult animals, and 4,745 from larvae; **Supplementary Figure 1C**). We reasoned that, besides technical variability (*e*.*g*. number of genes and UMIs per cell), cell heterogeneity in this dataset would depend on several biological variables, including the cell cycle stage, and the life stage (temporal axis) and pallial region (spatial or medio-lateral axis) from which each cell was sampled. Consistently, a UMAP embedding of RGCs/EGCs alone showed a clear separation between larval and adult cells (**Supplementary Figure 1C**). In previous work (Deryckere et al., 2025), we showed that the spatial axis in larvae is defined by gene expression gradients, and can be identified after regressing from the data the effect of other known sources of technical and biological variation (**Supplementary Figure 1D-E**). With the same approach, here we identified the spatial axis by fitting in the principal component (PCA) space a principal curve after regression of known sources of variance, and then used the medial marker *Wnt8b* to define the starting point of this principal curve (**Supplementary 1F-H**). Validating this approach, well-known and novel ventrolateral genes such as *Sfrp1* and *Sp5* and dorso-medial genes such as *Wnt8b* and *Foxp2* were expressed at opposite ends of this spatial axis curve (**Supplementary Figure 1I**). Progenitor cells were then assigned to their respective pallial region (MP, DP, LP, VP and AMY; **Supplementary Figure 1H, J**).

In the course of this analysis, we noticed that the genes with top loadings in PC1 and PC2 were cell cycle (PC1) and cellular quiescence genes (PC2). Quiescence is defined as a state of reversible cell cycle arrest (Morizet et al., 2025; Otsuki & Brand, 2018). We thus computed a new principal curve onto PC1 and PC2 to order cells along the gradient from deep quiescence to active proliferation, and identified 1,971 genes differentially expressed along this gradient. These genes included *Aqp4, Rarres1, Id1, Id2* and *Id3* (deep quiescence markers in mammalian NSCs (Urbán et al., 2019; Urbán & Cheung, 2021)), *Egfr, Ascl1* and *Foxo3* (mammalian shallow quiescence markers (Marqués-Torrejón et al., 2021; Urbán & Cheung, 2021)), and *Top2a, Pcna, Mcm2* and *Mcm7* (markers of actively proliferating cells, **Supplementary Figure 2A**). We then used hierarchical clustering (based on expression of proliferation and quiescence genes) to assign cells to 3 discrete groups: deep quiescence (dQ), shallow quiescence (sQ), and active (A) (**Figure 1E**; **Supplementary Figure 2A-B**). Notably, the adult pallium contained more progenitors in deep quiescence than in larvae (83% vs 38%, respectively, **Figure 1F**). Consistently, the deep quiescence cluster (dQ) was composed of adult EGCs (90.4% of total deep quiescent cells) and some cells from late larval stages, consistent with the notion that RGCs start exiting the cell cycle towards the end of larval development, before metamorphosis (Joven et al., 2018)(**Figure 1G**). Analysis of canonical quiescence and proliferation markers confirmed that across all pallial areas, deep quiescence genes were upregulated and active proliferation markers downregulated in adults compared to larvae (**Figure 1H**). The quiescence state differed not only across life stages but also across adult pallial regions: shallow quiescence genes were predominantly expressed in LP, while deep quiescence markers such as *Aqp4* were expressed in MP, DP and AMY (**Figure 1H**).

We validated these results with two complementary approaches. First, hybridization chain reaction (HCR) *in situ* hybridizations showed expression of *Gja1* and *Aqp4* (deep quiescence markers) in EGCs from all adult pallial regions except LP and VP, while *Sox4* (a marker of immature neurons) was predominantly expressed in LP and VP, suggesting higher rates of adult neurogenesis (**Figure 1I**). In contrast, in stage 55 larvae (the last larval stage before metamorphosis (Gallien & Durocher, 1957)) *Gja1* was confined to a few medial pallium EGCs, and *Sox4* was broadly expressed in the pallium (**Figure 1I**). Second, cumulative EdU administration in adult animals showed a higher EdU uptake in LP and VP EGCs, and low EdU labeling in other pallial regions, particularly in MP (**Supplementary Figure 2C-D**). *Pleurodeles* LP and VP EGCs may thus correspond to the proliferation hot spots identified in another newt species (Kirkham et al., 2014). Together, our results indicate profound differences of cell state between adult and larval neural stem cells, and across pallial regions in the adult. Most neural stem cells in larvae are actively proliferating, while in adults, most EGCs are quiescent, with regional variation in the depth of quiescence. These differences raise the question of whether the pallium responds to injury differently in larvae and adults. They also raise the more fundamental question of whether the neural stem cells states in adult newts are those more directly comparable to the stem cells and glial types in the adult mammalian brain.

### Deep quiescent EGCs in the adult newt brain express a mammalian astrocyte gene module

Salamander EGCs have been often compared to mammalian NSCs for their role in adult and regenerative neurogenesis. However, the sharp transcriptomic difference between newt RGCs and EGCs suggests that EGCs may arise from a true differentiation process, akin to the differentiation of astrocytes from RGCs during mammalian neural development. Clarifying what distinguishes salamander EGCs from mammalian astrocytes is important as it may reveal how astrocytes could be directed towards regenerative neurogenesis. We thus compared newt EGCs transcriptomic states to mammalian cell types. Specifically, we used an adult mouse dataset collected from the subventricular zone (SVZ), a neurogenic niche in the adult mouse brain. The dataset (Cebrian-Silla et al., 2025) included neural stem cells (B cells), intermediate progenitor cells (C cells), immature neurons called neuroblasts (A cells), and other glial cell types including parenchymal astrocytes (**Figure 2A**). We then analyzed the expression of genes that differentiate newt RGCs/EGCs states in this mouse dataset. For this, we computed differentially expressed genes (DEGs) in the newt dataset, and identified 130 genes upregulated in newt active neural stem cells, 177 genes in the shallow quiescence cluster and 126 genes in the deep quiescence cluster (**Figure 2B**). Then, we computed gene expression enrichment scores for these three gene sets in the mouse. The newt active gene module showed a strong enrichment in mouse dividing cells (**Figure 2C**). The newt shallow quiescence gene module was enriched in active B cells, C cells and dividing cells (**Figure 2D**). Lastly, the newt deep quiescence gene module showed high enrichment scores in astrocytes (**Figure 3E**). These analyses indicate that the deep quiescence state of newt EGCs, found in most adult pallial regions except the LP and VP, is defined by genes that distinguish astrocytes from neural stem cells in mammals.

**Figure 2.**
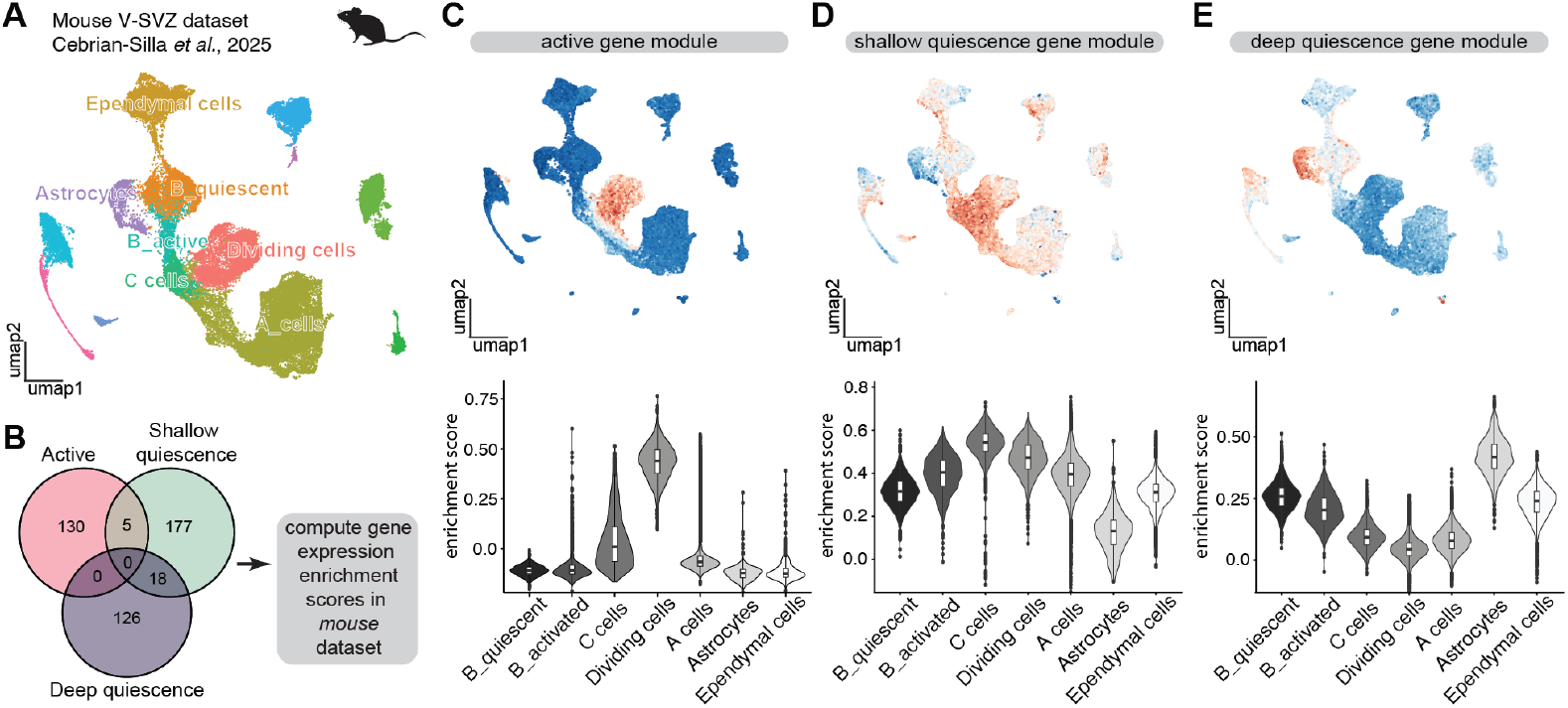
Deep quiescent ECGs in the newt pallium are defined by the expression of mammalian astrocytes genes. (A) UMAP plot of the Cebrian-Silla et al. 2025 mouse adult neurogenesis scRNAseq dataset. Cells colored by major class. (B) Venn diagram showing the intersection of the top 100 DEG that distinguish the three cell state clusters of newt neural stem cells. (C)-(E) Expression enrichment of newt cell state gene sets in the mouse adult neurogenesis data: newt active (C), shallow quiescence (D) and deep quiescence (E) gene sets. Top: UMAP plots of the mouse adult neurogenesis dataset with cells colored by expression enrichment. Bottom: violin plots showing expression enrichment scores in each mouse cell type.

**Figure 3.**
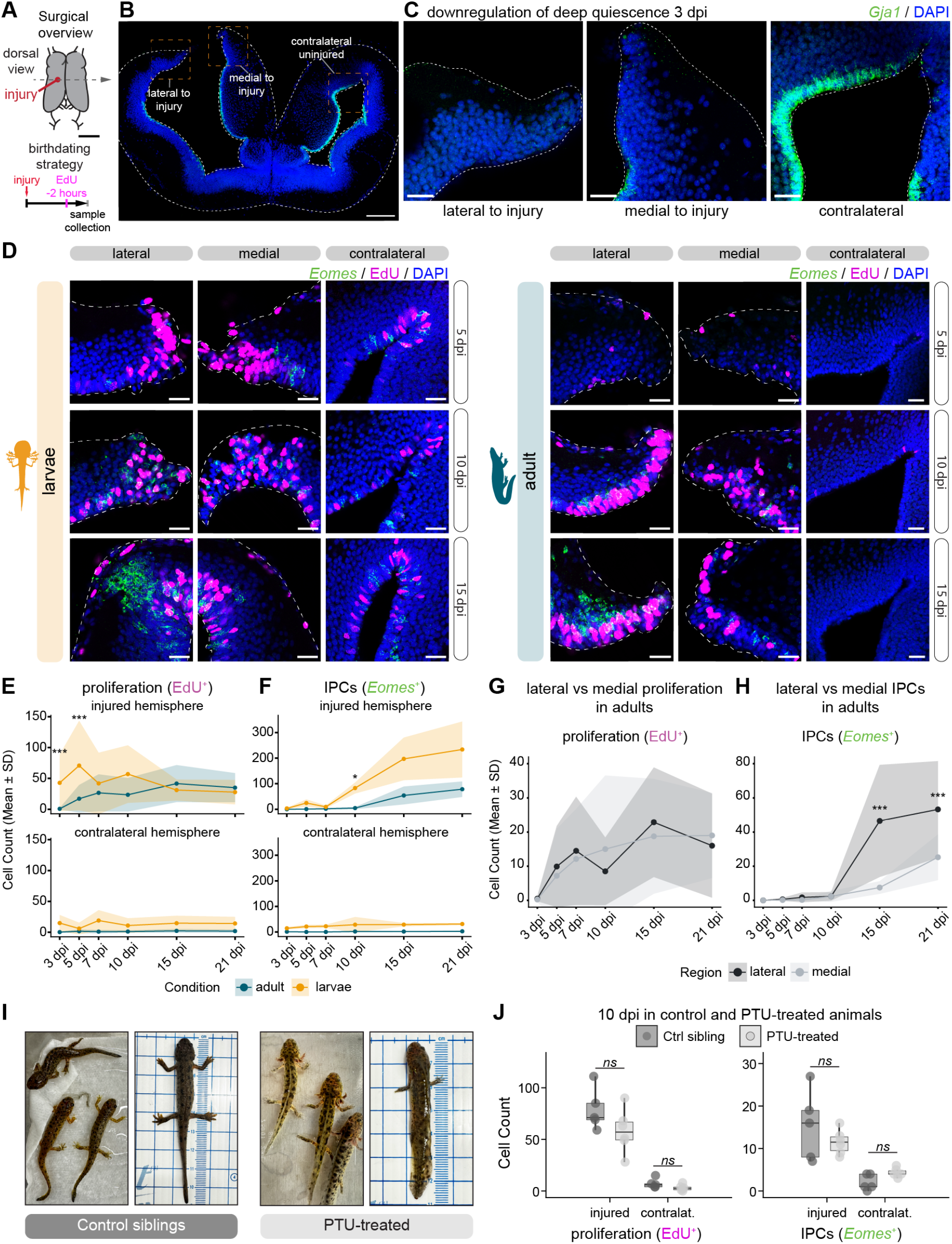
The regenerative response after brain injury is delayed in adult newts compared to larvae. (A) Schematic representation of the pulse-chase experiment and analysis. (B) Overview of a mid-telencephalic coronal section of an adult brain at 3 dpi, stained with the deep quiescent EGCs marker *Gja1*. Scale bar: 250 µm (C) Magnifications from (B) showing the downregulation of *Gja1* around the injury site when compared to the contralateral hemisphere. Scale bar: 50 µm. (D) Proliferating cells (EdU+, magenta) and IPCs (*Eomes*+, green) around the injury site and in the contralateral hemisphere, in larvae (left) and adults (right) at 5, 10, and 15 dpi. Images are closeups from coronal sections. Scale bar: 50 µm. (E) Mean cell counts of proliferating cells (EdU+) and (F) IPCs (*Eomes*+) from injured and contralateral hemispheres of larvae and adults. For each brain, cells were counted from three coronal sections, in a 200 µm-wide area centered on the injury site (see Methods). EdU+ and *Eomes*+ cells were quantified at 3, 5, 7, 10, 15, and 21 dpi. For each timepoint, n ≥ 3 adult animals and n > 5 larvae (see Methods). Ribbon limit represents standard deviation. Statistical significance was assessed using two-way ANOVA with post-hoc pairwise comparisons with Bonferroni correction. (G) Mean cell counts of proliferating cells (EdU+) and (H) IPCs (*Eomes+*) in the adult pallium, comparing lateral and medial regions relative to the injury site. Cells were counted from coronal sections as described above at 3, 5, 7, 10, 15, and 21 dpi. For each timepoint, n ≥ 3 adult animals (see Methods). Ribbon limit represents standard deviation. Statistical significance was assessed using two-way ANOVA with post-hoc pairwise comparisons with Bonferroni correction. (I) Images of PTU-treated newts and their control siblings, 188 days old, after 133 days of PTU treatment. (J) Quantification of proliferating cells (EdU+) and IPCs (*Eomes*+) at 10 dpi in PTU-treated animals and control siblings. Cell counting conducted as described above. All *p > 0*.*1*, Welch’s two-sample t-tests.

We next performed the reciprocal analysis, calculating DEGs of mouse cell types and analyzing expression enrichment in newt (**Supplementary Figure S3A**). The resulting modules contained 97 unique genes for active B cells, 92 unique genes for astrocytes and 95 unique genes for quiescent B cells (**Supplementary Figure S3A**). We observed expression of the mouse astrocyte gene module in all adult newt EGCs, but with LP EGCs showing the lowest astrocyte enrichment score (**Supplementary Figure S3B**). The quiescent B cell gene module followed a similar trend (**Supplementary Figure S3C**). Finally, when analyzing the active B cell gene module, we noticed a higher enrichment score in all larval RGCs on average compared to the adult EGCs (**Supplementary Figure S3D**). In the adult EGCs, however, we see that the highest scoring brain region was LP, followed by DP (**Supplementary Figure S3D**).

Taken together, our analysis indicates that newt neural stem cells exist in a continuum of molecular states — from deep quiescence to active progenitors — that are defined by the expression of genetic programs which in mammals define distinct cell identities. Importantly, the newt EGCs in deep quiescence express a battery of genes that differentiates mouse parenchymal astrocytes from neural stem cells. From an evolutionary perspective, these analyses suggest that newt EGCs differ from mammalian neural stem cells and astrocytes for their ability to flexibly access stemness and astrocytic programs in the same cell type.

### Different injury-response dynamics in late larvae and adults

The existence of EGCs in different quiescence states across life stages and pallial regions raises questions on how these quiescence states affect the regenerative response. To assess regeneration response in larvae and adults, we first developed a novel standardized pallial injury paradigm, consisting of a craniotomy and removal of the meninges followed by the excision of pallial tissue with a 250 µm-diameter biopsy punch (**Figure 3A**). This strategy allowed us to remove a reproducible-sized piece of tissue from the dorsal pallium and perform quantitative comparisons across individuals.

Our transcriptomic analysis predicts that, in the DP, adult EGCs need to transition from deep quiescence to an active state in order to proliferate in response to injury. To test this hypothesis, we performed surgeries in adults (*i*.*e*. post-metamorphic animals, over 13 months old) and then analyzed the regions immediately adjacent to the injury site, as well as the corresponding area in the contralateral uninjured hemisphere (**Figure 3B**). Previous work on axolotl demonstrated that the contralateral hemisphere can serve as an alternative control to sham surgeries (Amamoto et al., 2016). The expression of the deep quiescence markers *Aqp4* and *Gja1* in the dorsal pallium was downregulated around the injury site at 3 dpi (days post injury), suggesting that regenerative neurogenesis in the DP is preceded by the exit of EGCs from a deep quiescence state (**Figure 3B-C**; **Supplementary Figure S4A**).

We then asked whether the onset of injury-induced proliferation is different in larvae and adults, as adult EGCs need to exit quiescence and reenter the cell cycle, while a large proportion of larval RGCs are in active proliferation. We thus performed the same pallial injury paradigm on late-stage larvae (stage 55, ∼90 days post fertilization) alongside adults, and labeled S-phase cells at multiple time points (3, 5, 7, 10, 15 and 21 dpi) by administering the thymidine analog EdU 2 hours before brain collection (**Figure 3A**). Additionally, we labeled IPCs, which are neurally-committed progenitors that amplify the neuronal output of neural stem cells, by staining for *Eomes* (*Tbr2*) (**Figure 3D**, **Supplementary Figure S4B**). Across all conditions, we quantified the number of EdU+ and *Eomes*+ cells in the 100 µm-wide regions medial and lateral to the injury site and in a corresponding 200 µm-wide portion of the pallium in the contralateral hemisphere.

In late larvae, pallial injury enhanced ongoing proliferation, with a significantly higher number of EdU+ cells at the site of injury already at 3 dpi compared to the contralateral hemisphere (*p<0*.*05*; this and all other comparisons reported below are ANOVAs followed by Bonferroni-adjusted post-hoc pairwise comparison). Such response is sustained at 5, 7 and 10 dpi (*p<0*.*0001, p<0*.*05* and *p<0*.*001*, respectively) and then decreases at 15 and 21 dpi to similar levels found in the contralateral hemisphere. In adults, the onset of proliferation was delayed compared to larvae: there was almost no proliferation around the injury site at 3 dpi, and no differences compared to the contralateral hemisphere. However, from 5 dpi onward, the injury site exhibited significantly more EdU+ cells than the contralateral hemisphere (*p<0*.*0005* for all timepoints; **Figure 3E**). Moreover, our analysis showed that the number of EdU+ cells was significantly higher in larvae compared to adults (*p<0*.*0001*, two-way ANOVA), with the biggest differences observed at 3 and 5 dpi (**Figure 3E**). Taken together, our results indicate that adults have a later onset of proliferation and a lower number of EdU+ cells around the injury site when compared to larvae. We also found significant differences for IPCs. In larvae, the number of *Eomes*+ cells showed a steep and significant increase between day 7 and 10 post injury (*p<0*.*05*) and then remained elevated at subsequent timepoints compared to the contralateral hemisphere (*p<0*.*0001* for both 15 and 21 dpi), suggesting an injury-induced increase of neurogenic divisions from 7 dpi onwards (**Figure 3D**, **Supplementary Figure S4B**). However, in adults, the number of *Eomes*+ cells started to increase only between 10 and 15 dpi and remained elevated at 21 dpi compared to the contralateral hemisphere (**Figure 3D**), showing that the onset of IPCs generation is also delayed in adults compared to late larvae (**Figure 3F**). These results indicate that the dynamics of neural stem cell proliferation and of regenerative neurogenesis differ in larvae and adults, possibly as a result of the different quiescence states of RGCs and EGCs at the time of injury.

We then asked whether, in the adult, there would be any difference between the injury response of EGCs at the medial side of the injury in DP, where EGCs are in deep quiescence, and at the lateral side of the injury, which includes shallow quiescent EGCs in LP. While the number of proliferating cells was not substantially different on the two sides (**Figure 3G**), we found a higher number of *Eomes*+ IPCs lateral to the injury side compared to the medial side of the injury at 15 dpi and 21 dpi (**Figure 3H**, **Supplementary Figure S4B**). This suggests that in adults the EGCs lateral to the injury side give rise to a larger number of neurons through IPC-mediated indirect neurogenesis, and thus may have a disproportionate contribution to regeneration.

Studies in other regenerative systems have pointed out how metamorphosis coincides with a decline of regenerative capacity (Monaghan et al., 2014). We thus wondered whether the different dynamics of the regenerative response in larvae and adults were solely accounted for by metamorphosis. To test this, we treated a cohort of stage 54 larvae (79 days old) with propylthiouracyl (PTU), an established thyroid hormone antagonist that suppresses metamorphosis in anamniotes (Degitz et al., 2005; Sperry & Grobstein, 1985). A cohort of sibling animals was housed in regular tank water as a control. Animals underwent PTU treatment for 51 weeks before the injury surgery. PTU-treated animals retained a pre-metamorphic phenotype for the entire duration of the treatment, as evidenced by the retention of gills and of a wide aquatic tail, while control siblings went through metamorphosis starting at 3 months after fertilization (**Figure 3I**). We then measured EdU uptake and *Eomes* expression in both cohorts 10 days after pallial injury. At this time point, *Eomes* is upregulated in larvae but not in adults; therefore, we expected to see more *Eomes+* cells in PTU-treated animals compared to the controls, under the assumption that PTU animals would exhibit a larval regenerative response (**Figure 3E-F**). Surprisingly, we did not observe any significant difference between PTU-treated and control animals in the number of EdU+ cells or IPCs (**Figure 3J**). Indeed, EGCs deep quiescence markers were expressed with identical patterns in PTU-treated animals and in their metamorphosed siblings (**Supplementary Figure S3C**). These results suggest that the developmental progression of EGCs towards quiescence is not simply the consequence of metamorphosis but might be partly associated with aging. Together, our data indicates that the molecular state of EGCs biases not only the rate of homeostatic neurogenesis across life stages (Joven et al., 2018), but also the onset and progression of the regenerative response after injury.

### *Pleurodeles waltl* regenerates neurons in the pallium after a mechanical injury

The differences in early response to pallial injury in larvae and adults made us wonder when regenerative neurogenesis is completed, and whether the final outcomes of regeneration at the cell type level differ across life stages.

To determine an endpoint for analysis, we performed the aforementioned pallial injury paradigm (**Figure 3A**). In the adult, wound closure was visible around 21 dpi; the injured hemisphere looked indistinguishable from the contralateral side at 84 dpi (**Figure 4A**). To label neurons born after the injury, we administered EdU daily for five consecutive days, from 5 dpi until 9 dpi (**Figure 4B**), given that proliferation in adults is not upregulated until 5 dpi (**Figure 3E**). Whole mount light-sheet imaging of cleared regenerated brains showed a spot with high density of EdU labeling in the pallium, corresponding to the regenerated area, and an overall increase of EdU labeling on the surface of the injured hemisphere, as a result of the regeneration of the meninges (**Figure 4C**).

**Figure 4.**
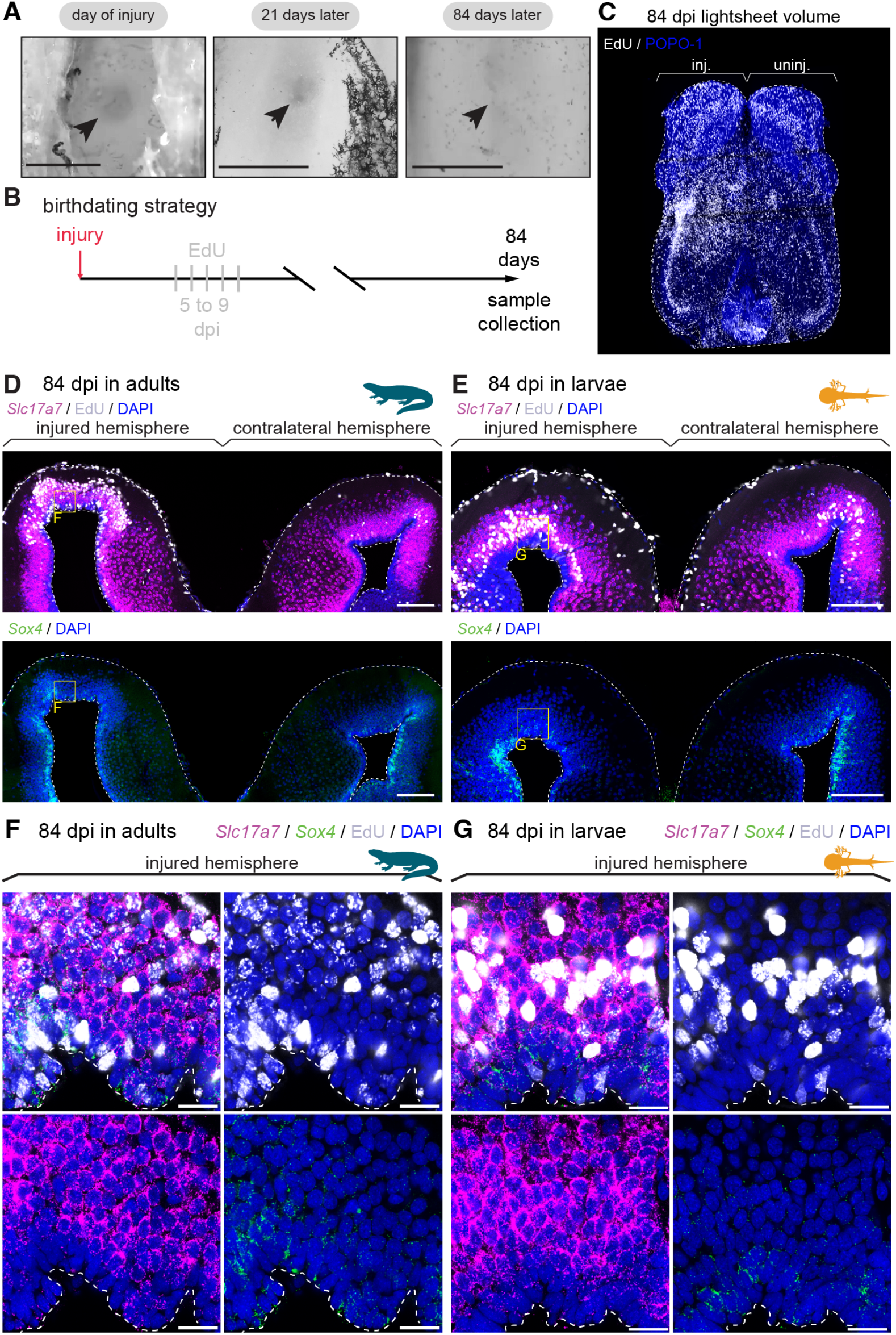
Regenerative neurogenesis of the newt pallium is complete by 84 days post-injury. (A) Dorsal view of representative craniotomies and pallial injuries at 0, 21, and 84 days post injury (dpi). The hole produced by the biopsy punch is visible at 0 and 21 dpi but not at 84 dpi. Scale bar 500 µm. (B) EdU pulse-chase strategy to label neurons born between 5 and 9 dpi. (C) Dorsal view of whole-mount staining of EdU-labeled cells (white) highlighting the injury site (left hemisphere) in a brain collected at 84 dpi and processed with clearing and light sheet imaging. White dashed lines depict borders of the telencephalon. inj.: injured hemisphere. uninj.: contralateral hemisphere. (D)-(E) Coronal sections showing pallial hemispheres in an adult (D) and a late larva (E) at 84 dpi. A high-density of EdU+ cells (white) labels the injury site. Mature glutamatergic neurons and immature neurons are labeled by *Slc17a7* (Vglut2, magenta) and *Sox4* (green) expression, respectively. Scale bars: 200 µm. (F)-(G) Magnifications of the regenerated dorsal pallium from (D)-(E), showing colocalization of EdU and *Slc17a7* and a low number of *Sox4+* cells. Scale bars: 30 µm.

To test whether regenerative neurogenesis was still ongoing or had already ended at 84 dpi in adults, we stained the regenerated pallium at 84 dpi with *Slc17a7* (Vglut2, marker of differentiated glutamatergic neurons) and *Sox4* (marker of immature neurons). In the injured hemisphere, most of the cells born between 5 and 9 dpi were mature glutamatergic neurons (*Slc17a7*+ EdU+) that had integrated in the neuronal layers. We also found a few EdU+ EGCs in the ventricular zone (**Figure 4D, 4F**; EdU in white). Notably, there were few *Sox4+* immature neurons in the injured DP close to the ventricular surface, with a pattern comparable to the contralateral (uninjured) hemisphere, suggesting that by 84 dpi, regenerative neurogenesis in the DP had terminated (**Figure 4D, 4F**). Next, we performed the same experiment in stage 55 larvae. All of the injured larvae underwent metamorphosis between injury and brain collection at 84 dpi. Similar to the adults, there were few *Sox4+* immature neurons remaining in the injured hemisphere, indicating the conclusion of regenerative neurogenesis by 84 dpi (**Figure 4E, 4G**). Together, these results show that our injury paradigm is effective and that regenerative neurogenesis, as measured by return of immature neurons to baseline, is complete by 84 dpi at the histological level.

### Spatial patterning is not restored correctly in the regenerated adult pallium

The newt pallium is organized along the medio-lateral axis into regions defined by the spatially-restricted distribution of glutamatergic neuron types (Woych et al., 2022). Experiments in axolotl indicate that regional boundaries are largely reestablished after a pallial injury (Amamoto et al., 2016; Wei et al., 2022). However, our analysis of the early response to pallial injury in the newt (**Figure 3**) suggested that EGCs lateral to the injury, which include EGCs in the LP, produce more IPCs and therefore may have a disproportionate contribution to regeneration. To test this hypothesis, we analyzed the molecular identity of the neuron types produced after a DP injury. Pallial injuries on late-stage larvae (stage 55) and adults were followed by EdU injections between 5 and 9 dpi (**Figure 5A**). We then analyzed regenerated brains at 84 dpi and selected three marker genes that together label all major cell classes in the DP and LP: *Nts* for DP superficial layer neurons (DP-SL) and parts of the VP, *Satb1* for LP glutamatergic neurons, and *Sox2* for EGCs (periventricular) and GABAergic interneurons (parenchymal, including *Satb1*+ and *Nts*+ subsets) (**Figure 5B**). We thus counted the total number of each molecularly identified cell type in the entire DP and LP from three consecutive 70 µm-thick coronal sections, spanning approximately 210 µm (along the rostrocaudal axis) centered on the injury site. We also counted cells in the contralateral hemisphere. The analysis of regional boundaries is described in **Figure 5**, while the regeneration of DP-SL and DP deep layer neurons (DP-DL) is shown in **Figure7**.

**Figure 5.**
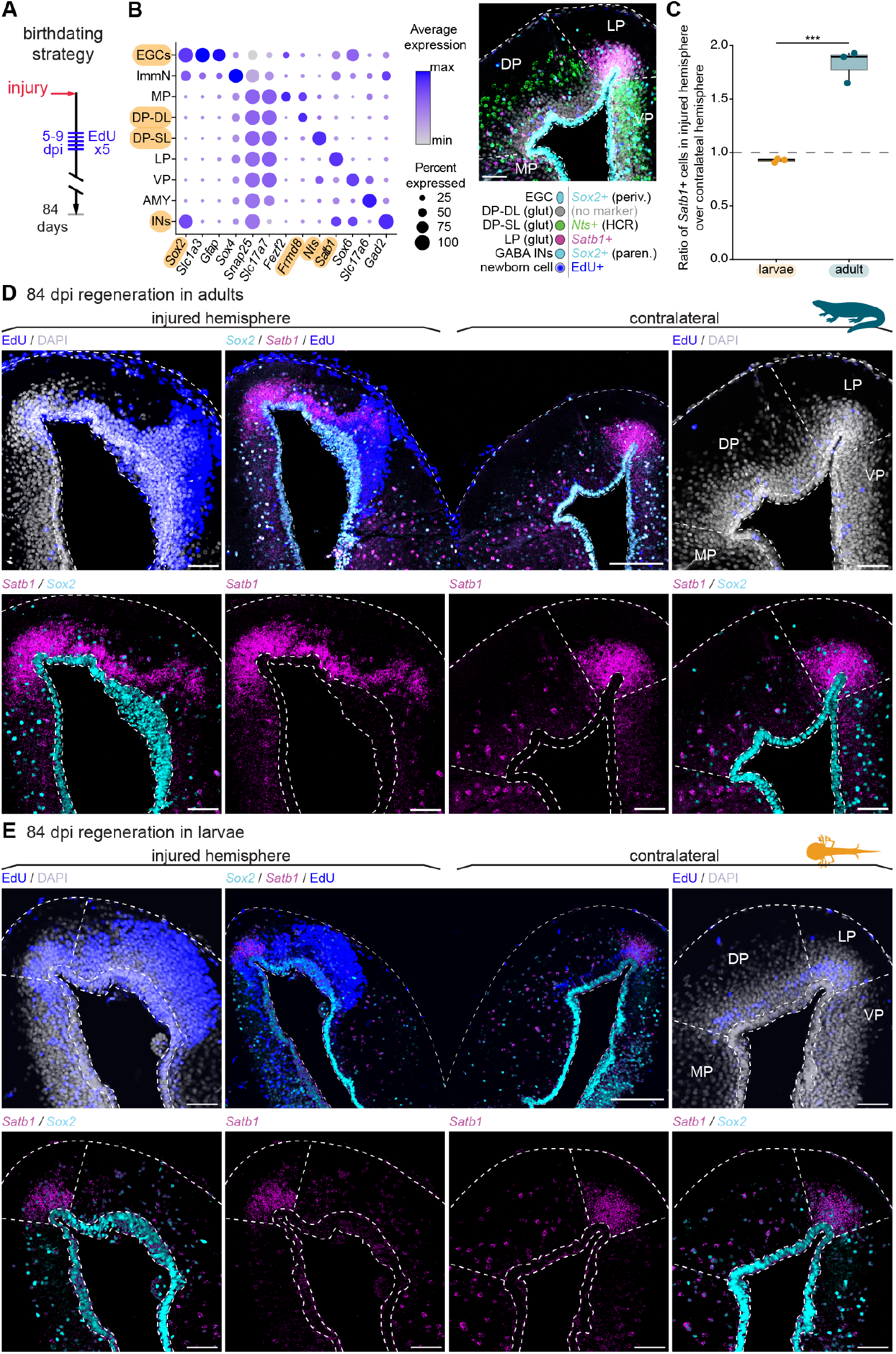
Pallial injuries regenerate with an overrepresentation of lateral pallium *Satb1*+ neurons in adult newts but not in larvae. EdU pulse-chase labeling of cells born between 5 and 9 dpi. (B) Left: marker gene expression in pallial EGCs and glutamatergic neuron types. Dot size indicates percentage of cells expressing the gene, color indicates expression level. Right: Coronal section of the pallium stained with the panel of selected markers: EdU (blue), Sox2 immunohistochemistry (cyan), and *Nts* and *Satb1* HCRs (green and magenta, respectively). Scale bar: 50 µm. (C) Ratio between the total number of LP *Satb1*+ neurons in the injured hemispheres and the total number of LP *Satb1*+ neurons in the contralateral hemisphere, in adult and larval regenerates. Cells were counted in 3 sections spanning the injury site (see Methods). *p = 0*.*009*, Welch’s two-sample t-test, n = 3 per condition (D)-(E) Coronal sections through the pallium 84 days after injury in adult (D) and larval (E) newts. Injury site labeled by EdU (blue); Sox2 and *Satb1* expression shown in cyan and magenta, respectively. Scale bars: 250 µm in overview sections and 100 µm in insets.

Quantifications showed that after regeneration in the adult animals, the number of LP *Satb1*+ cells in the injured hemisphere was nearly double the number of LP *Satb1*+ cells in the contralateral hemisphere (**Figure 5C**). This increase of LP *Satb1*+ cells was caused by the generation of new *Satb1*+ cells after injury, as ∼30% of all EdU-labeled glutamatergic neurons were *Satb1+* (**Supplementary Table 1**). Conversely, in larvae, the total number of LP *Satb1*+ cells was approximately the same on the injured and contralateral sides (**Figure 5C**). Consistently, in adult animals, the LP domain labelled by *Satb1* in the injured hemisphere extended towards the medial side of the pallium, invading the DP territory (**Figure 5D**, EdU in blue). In contrast, in animals injured as larvae, *Satb1+* cells remained confined to the LP territory, and the boundary between LP and DP was well preserved (**Figure 5E**).

The different regeneration outcomes in larvae and adults raise two alternative hypotheses. Neural progenitors at these two life stages may respond differently to spatial patterning cues that reorganize the regenerating pallium along the medio-lateral axis. In this case, in the adult, neural progenitors would be responding to lateral but not medial patterning cues, with LP *Satb1*+ cells forming at the expense of DP neurons. Alternatively, neural progenitors could maintain the regional identity established during development, but with LP progenitors producing more neurons than DP progenitors during regeneration in the adult. In this case, we would expect to see that the new LP *Satb1*+ neurons are produced in addition to, and not at the expense of DP neurons. Quantifications supported the latter scenario: in the adult, but not in larvae, the total number of glutamatergic neurons across the DP and LP was on average ∼26% larger in the injured hemisphere compared to the contralateral hemisphere (**Supplementary Figure S5A-B**). We thus propose that the EGCs participating in pallial regeneration have distinct and fixed regional identities, and that their relative contribution to the regenerated tissue depends on their initial quiescence state, which changes throughout the animal life cycle. The higher neurogenic output of LP progenitors, which enter regeneration from a shallow quiescence state, may thus underlie the production of a larger number of IPCs (**Figure 3**) and expansion of LP cells after an injury of the DP.

### Adult neurogenesis in the newt dorsal pallium is developmentally constrained

Given that spatial patterning is disrupted in the regenerating adult pallium, we next asked how the regenerating adult brain would compare to homeostatic conditions in terms of *temporal* patterning. In the developing DP, temporal patterning of multipotent RGCs produces an early-born superficial layer (DP-SL) and a late-born deep layer (DP-DL) (Deryckere et al., 2025) (**Figure6A**). However, it is unclear whether both DP early-born *Nts*+ and late-born *Frmd8*+ neurons are produced after metamorphosis during adult neurogenesis, or if the output of adult EGCs is restricted to late-born *Frmd8*+ neurons. To answer this question, we performed EdU pulse-chase experiments on adult *Pleurodeles* (age: 12.4 months, EdU injected for 5 consecutive days). EdU+ labeled cells were analyzed 84 or 180 days after injection (dai), together with established cell type markers: *Sox2*, which labels EGCs (elongated, adjacent to the ventricle) and interneurons (round, parenchymal), *Frmd8*, and *Nts* (**Figure 5B**). We quantified the total number of EdU+ cells in the DP and LP of the adult brain, from sections spanning approximately 550 µm along the rostrocaudal axis (n=3 animals for each time point, 8 sections, 1106 total EdU+ cells). We detected a modest number of EdU+ EGCs (**Figure6B-C**), consistent with the idea that the neural stem cell pool is maintained through asymmetric divisions. The number of EdU+ *Sox2*+ parenchymal cells, which are newly generated pallial interneurons, was negligible. In the LP (demarcated by lack of *Nts* and *Frmd8* expression; **Figure 5B**), we found abundant EdU+ neurons (**Figure6B-C**), indicating that this region has a relatively high rate of adult neurogenesis. In the DP, we found EdU+ *Frmd8*+ cells, indicating that DP-DL neurons are produced during adult neurogenesis. However, we did not detect a single newborn DP-SL neuron (EdU+ *Nts*+) in the screened region in any of the conditions assessed (**Figure6B-C**). These results indicate that adult newts do not generate early-born DP-SL neurons under homeostatic conditions, and that pallial EGCs retain the temporal fate restriction put into place during development.

**Figure 6.**
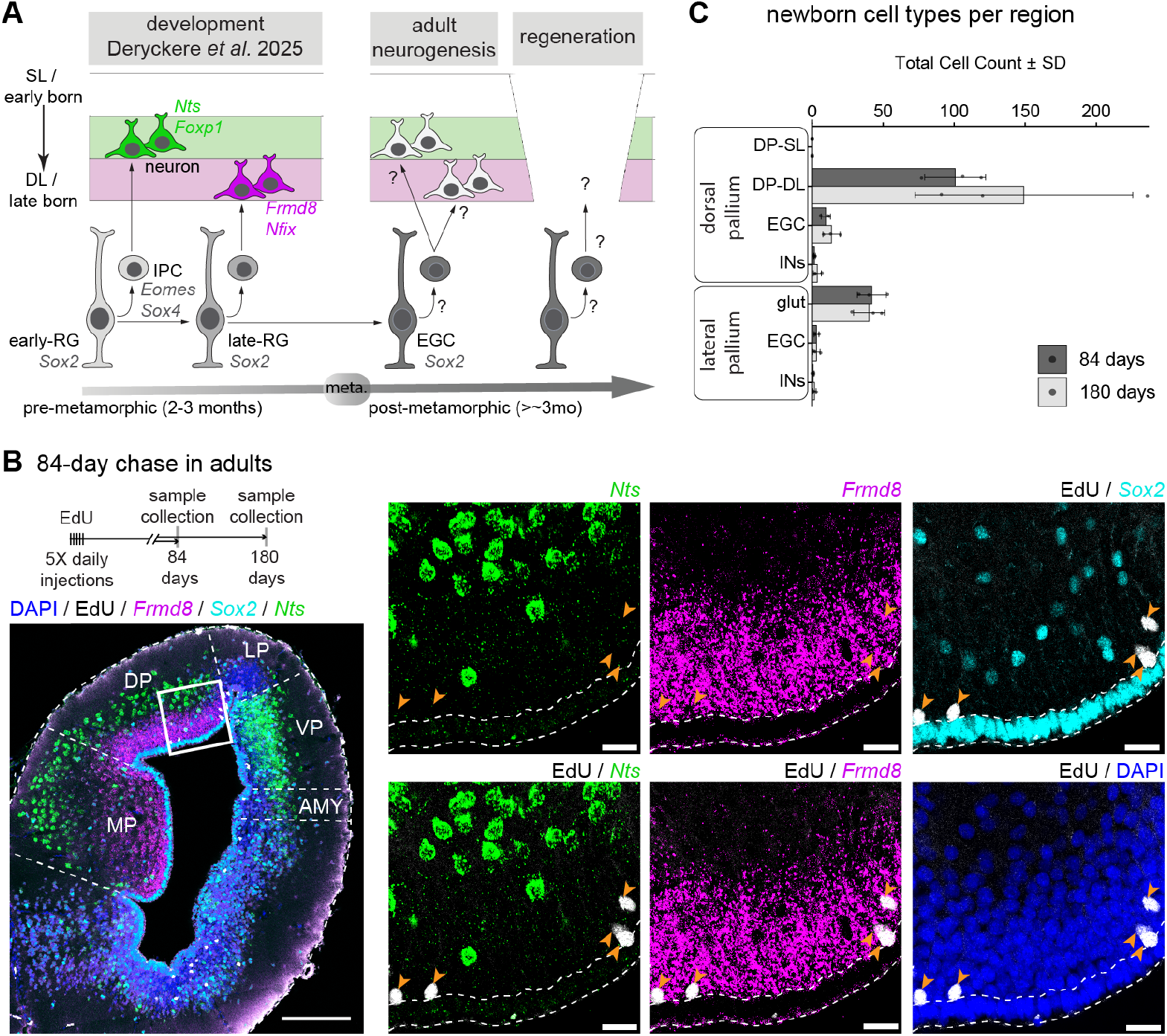
Homeostatic adult neurogenesis in the newt DP is limited to late-born deep layer neurons. (A) Schematic representation of DP development and open questions. (B) Top left: EdU pulse-chase strategy for assessing adult neurogenesis. Bottom left: Coronal section at mid-telencephalic level showing EdU-labeled cells (white), Sox2 antibody staining (cyan), and *Nts* and *Frmd8* HCR staining (green and magenta, respectively) at 84 days after EdU injection. Right: magnifications of the dorsal pallium, arrowheads indicate EdU+ neurons. Scale bar: 250 µm in the overview image and 30 µm in the insets. (C) Cell type identity of EdU+ cells in the DP and LP, based on morphological landmarks and maker expression (see also Figure 5B). Cells were counted across 8 consecutive coronal sections, spanning 560 µm in total along the rostrocaudal axis. n = 3 animals for each time point, error bars represent standard deviation.

### Regeneration of early-born neurons in the adult dorsal pallium

Under the hypothesis that regenerative neurogenesis would follow the same temporal constraints of adult neurogenesis, we expected to see only newly-born DP-DL neurons in the regenerated pallium in both late-stage larvae (stage 55, after the stage 50 switch from DP-SL to DP-DL, **Figure6A**) and adults. To test this, we analyzed the molecular identity of neurons in the regenerated DP (analysis at 84 dpi with EdU administration between 5 and 9 dpi). Indeed, the vast majority of the EdU+ cells in the DP expressed the deep layer marker *Frmd8* (**Figure7A**).

**Figure 7.**
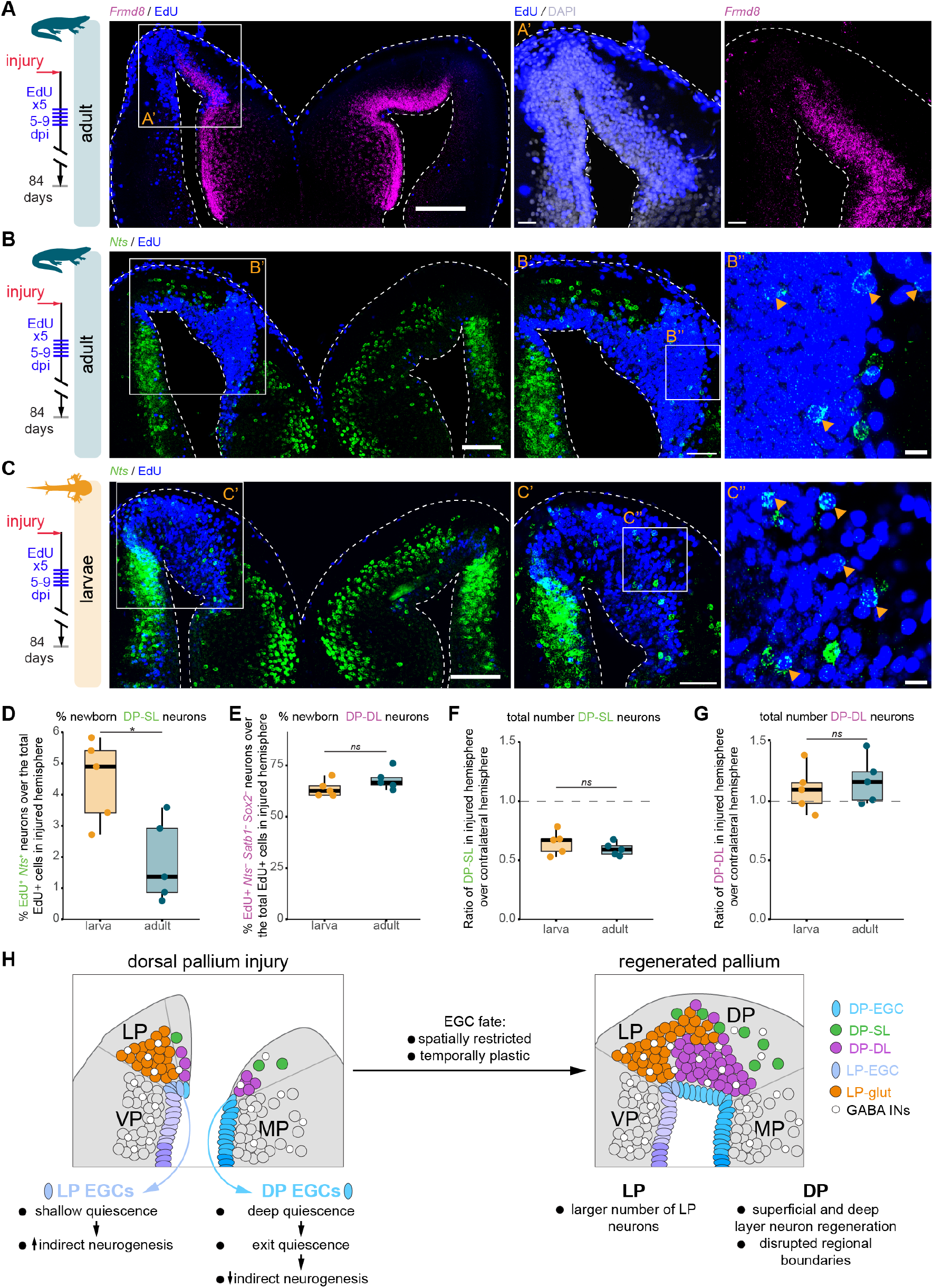
EGCs override temporal patterning restrictions and regenerate both early- and late-born DP neurons. (A)-(A’’) Overview (A) and magnifications (A’-A’’) of a coronal section from an adult regenerated pallium at 84 dpi, after EdU pulse-chase injections between 5 and 9 dpi. EdU (blue) and the DP-DL marker *Frmd8* (HCR, magenta) are colocalized, indicating regeneration of DP-DL neurons. Scale bars: 250 µm (A) and 50 µm (A’-A’’). (B)-(B’’) Overview (B) and magnifications (B’-B’’) of a coronal section from an adult regenerated pallium at 84 dpi, after EdU injections between 5 and 9 dpi. Some EdU+ cells (blue) coexpress the DP-SL marker with *Nts* (HCR, green), indicating regeneration of DP-DL neurons. Scale bars: 250 µm (B), 50 µm (B’) and 20 µm (B’’). (C)-(C’’) As in (B)-(B’’), but after an injury of a larval brain. Scale bars: 200 µm (C), 100 µm (C’) and 20 µm (C’’). (D) Percentage of EdU+ *Nts*+ cells over the total number of EdU+ cells in the regenerated pallium (DP and LP) after adult and larval injuries. *p = 0*.*015*, Welch’s two-sample t-test, n = 5 animals per condition. (E) Percentage of EdU+ DP deep-layer cells (*i*.*e*. EdU+ *Nts-Satb1-* neurons, which are putative *Frmd8*+ neurons) over the total number of EdU+ cells in the regenerated pallium (DP and LP) after adult and larval injuries. *p = 0*.*17*, Welch’s two-sample t-test. (F) Ratio of total number of *Nts*+ cells in the DP of the injured hemisphere relative to the total number in the contralateral DP; comparison of larvae and adults. *p = 0*.*338*, Welch’s two-sample t-test. n = 5 animals per condition (G) Ratio of total number of DP-DL (*Nts-Satb1-*) neurons in the DP of the injured hemisphere relative to the total number in the contralateral DP; comparison of larvae and adults. *p = 0*.*575*, Welch’s two-sample t-test. n = 5 animals per condition. (H) Model illustrating how the constraints on EGCs spatial patterning and plasticity of temporal patterning would produce the observed regeneration phenotypes in adult newts.

After staining for the DP-SL marker *Nts*, we observed that the *Nts*+ layer was dramatically disrupted in the injured hemisphere compared to the contralateral hemisphere in both adults and larvae (**Figure7B-C**). Surprisingly, we could still identify some EdU+ *Nts*+ neurons in both stages (**Figure7B’’-C’’**). Importantly, we did not observe any EdU+ *Nts*+ cells on the contralateral uninjured side, consistent with our adult neurogenesis data (**Figure7B-C**). After quantifications, we found that in adults, on average 1.9% of the total EdU+ cells in the regenerating pallium were also positive for *Nts*. In contrast, in larvae, 4.5% of all EdU+ cells were *Nts*+ (**Figure7D**). The proportion of EdU+ cells that were DP-DL neurons in the regenerating pallium were similar for larvae and adults (**Figure7E**). These findings mark a clear difference between regenerative and adult neurogenesis (**Figure6**). While in homeostatic conditions, the neuronal output of EGCs is limited to late-born DP-DL neurons, pallial injury induces the generation of neurons that would normally only be produced early in development.

We next wondered whether the different proportions of EdU+ *Nts*+ neurons in larvae and adults indicated a better capacity of late larvae to faithfully regenerate the original cellular diversity. Remarkably, quantification of the total number of DP-SL neurons in the injured and contralateral hemisphere revealed that the number of DP-SL neurons in the regenerated hemisphere was consistently lower compared to the contralateral hemisphere in both life stages, indicating that DP-SL neurons were not restored back to baseline (**Figure7F**). Conversely, the total number of DP-DL neurons was slightly higher in the regenerated hemisphere compared to the contralateral one, again in both larvae and adults (**Figure7G**). Our results thus indicate that larvae regenerated DP-SL neurons earlier than adults, because although the relative proportions of DP-SL neurons after regeneration was similar in the two life stages (**Figure7F-G**), larvae had a higher proportion of EdU+ *Nts*+ cells labeled between 5 and 9 dpi (**Figure7D-E**). This observation is consistent with our analysis at early time points after pallial injury, where the onset of proliferation in adults was delayed in comparison to larvae (**Figure 3**).

Taken together, our results show that regenerative neurogenesis is not simply an amplification of adult neurogenesis but rather a unique state, where EGCs in the DP override restrictions imposed by temporal patterning and gain access to early neuronal differentiation programs. We thus conclude that newt DP neural stem cells regenerate the original cell type repertoire lost to injury, although not in perfect proportions, regardless of whether they are active radial glia progenitors in larvae or astrocyte-like deep quiescent ependymoglia in adults.

## Discussion

Why regeneration is restricted to a limited range of vertebrate species remains a mystery. Several hypotheses have been proposed to explain why the capacity to regenerate the central nervous system was seemingly lost multiple times in vertebrate evolution (Alibardi, 2023; Blackshaw, 2022; Tanaka & Ferretti, 2009). However, these hypotheses have remained largely untested, given that CNS regeneration has been studied in a limited number of species. Here, we characterized brain regeneration in the Iberian ribbed newt *Pleurodeles waltl*, a salamander species that is separated from the axolotl by ∼150 million years, and has a complete life cycle including a true post-metamorphic stage. Our results reveal that the regenerative capacity of newts is age-dependent, and modulated by the availability of a diverse set of neural stem cells in varying spatial and temporal molecular states. When analyzed in a broader comparative framework, these findings challenge current hypotheses on the nature and evolution of brain regeneration.

In vertebrates, brain regeneration correlates with the abundance of neural stem cells and presence of adult neurogenesis. Studies in several regenerative models, including newts and axolotl, have shown that ependymoglia cells act as neural stem cells and produce new neurons in response to brain injuries (Berg et al., 2011; Fu et al., 2026; Goldshmit et al., 2012; Hui et al., 2010; Kaslin et al., 2013; Maden et al., 2013; Mchedlishvili et al., 2012; Parish et al., 2007; Rodrigo Albors et al., 2015). These observations support the hypothesis that regeneration has been lost as the indirect consequence of the loss of neural stem cells and continuous tissue growth, including adult neurogenesis (Blackshaw, 2022). A corollary of this hypothesis is that regenerative neurogenesis is an amplified version of adult neurogenesis. Instead, our data clearly show that regenerative neurogenesis in the newt pallium operates within a different set of constraints than those which govern adult neurogenesis. Building on detailed knowledge on pallial neuron types (Woych et al., 2022) and their development (Deryckere et al., 2025), we found that adult neurogenesis in the dorsal pallium is restricted to late-born neurons, while early-born neurons are not produced in homeostatic conditions. This developmental restriction is overturned in response to brain injury, not only in post-metamorphic newts, but also in larvae that have already grown past the switch from early to late neurogenesis. These observations indicate that brain regeneration in newts is not simply explained by the presence of neural stem cells and adult neurogenesis, but rather suggest the existence of regeneration-specific molecular pathways that can recruit differentiation programs normally active only during early development. Supporting this hypothesis, a regeneration-specific ependymoglia type has been recently discovered in the axolotl brain (Lust et al., 2022; Wei et al., 2022). Failure to activate this regeneration-specific program may explain, at least in part, the limited brain regenerative capacity of reptiles, where ependymoglia cells express many of the same stemness markers found in salamander ependymoglia (Clinton et al., 2014; Tosches et al., 2018; Trujillo-Cenóz et al., 2014).

Across vertebrates, regenerative capacity correlates with levels of thyroid hormones (Hirose et al., 2019). In amphibians, regeneration typically declines after metamorphosis, which coincides with an increase in thyroid hormone signalling (Degitz et al., 2005; Frangioni et al., 2006; Smirnov et al., 2020; Sperry & Grobstein, 1985). These observations lend support to the hypothesis that regeneration was lost in land vertebrates as a consequence of an increase in thyroid hormone signaling and overall metabolic rates (Alibardi, 2023). However, the effects of thyroid hormones on neural stem cells biology are complex (Valcárcel-Hernández et al., 2024). Thyroid hormone signaling seems to be primarily related to the control of energy expenditure during environmental transitions, with the effects on regeneration being potentially secondary and context-dependent (Zwahlen et al., 2024). Here, studying regeneration in a salamander with a complete life cycle provided the opportunity to compare pre- and post-metamorphic stages. Our results show that, while the regenerative response is indeed slower in older animals, the same cell types regenerate in both life cycle stages, consistent with recent results on lens regeneration (Tsissios et al., 2023). Moreover, suppression of thyroid hormone signaling with PTU was not sufficient to accelerate regeneration to a larval pace, suggesting that regeneration is not modulated by a discrete metamorphic switch, but rather by aging.

In the axolotl, which does not naturally undergo metamorphosis, the relationship between aging and brain regeneration has been poorly explored. The transcriptomic cell-type analyses of pallial regeneration conducted in young axolotls are largely consistent with our newt larval data (Lust et al., 2022; Wei et al., 2022). However, pallial lesions in older axolotls produced some morphological defects resembling our adult newt phenotypes, including medial displacement of LP neurons (Amamoto et al., 2016). Further comparisons of regeneration in young and older axolotls could help disentangle the relative influence of metamorphosis and aging on the regeneration process.

Our analysis of molecular states in NSCs provides a key framework to understand the regeneration phenotypes we describe in the newt pallium. The most dramatic difference we identified – the presence of an aberrant proportion of LP *Satb1*+ cells in adults – could be fully explained by the different molecular states found in the adult EGCs participating in regeneration. Only in adults, but not in larvae, DP EGCs are in a deep quiescence state, have a low proliferative rate, and need to downregulate deep quiescence markers before initiating a neurogenic response. Conversely, LP EGCs are active or in shallow quiescence, have a higher neuronal output through IPCs, and contribute a larger number of LP *Satb1*+ neurons to the regenerated tissue. Our model assumes that the pallium is not spatially repatterned after an injury but the fate of EGCs remains regionally restricted. At the endpoint of regeneration, the proportion of different neuron types is then simply determined by the relative contributions of LP and DP progenitors (**Figure7H**). Our results thus suggest that, while temporal patterning is more plastic such that temporal restrictions can be reversed after an injury, spatial patterning defines pools of progenitors that remain lineage-restricted in every condition. Brain regeneration is thus substantially different from regeneration in other systems such as the limb, where the blastema is repatterned to recreate all parts of the limb in the correct proportions (Otsuki & Tanaka, 2022).

Interestingly, the transcriptome of deep quiescent NSCs stands out for the expression of a mammalian astrocyte gene program. The downregulation of deep quiescence markers in response to injury could thus be described as dedifferentiation, analogous to the injury-induced dedifferentiation of ependymoglia cells in the axolotl spinal cord (Arsanto et al., 1992; Echeverri & Tanaka, 2002) and of Müller glia cells in the zebrafish retina (Lyu et al., 2023). Similarly, these glial cell types also share the capacity to re-access early differentiation programs during regeneration (Lyu et al., 2023). Plasticity at the gene regulatory level may thus underlie CNS regeneration across regions and species. This plasticity may have been lost in mammals, where the genetic programs that define adult NSCs and parenchymal astrocytes remain segregated in distinct and specialized cell types (Laywell et al., 2000; Tiwari et al., 2018; Velloso et al., 2022). An irreversible segregation of gene regulatory programs across cell types in mammals may explain why their astrocytes respond to brain injury with reactive gliosis instead of neurogenesis. We thus propose that a continuum of cell states, ranging from a deep quiescence astrocyte-like state to an active neural stem cell state, provides newt ependymoglia cells the capacity to operate as astrocytes while also serving as neural stem cells. A cellular division of labor in mammalian ancestors, with the evolution of distinct astrocytes and neuronal stem cells, may have favored further functional specializations at the cost of cellular plasticity, and ultimately, of regenerative capacity. With its growing experimental toolkit (Elewa et al., 2017; Hayashi et al., 2013; Jaeger et al., 2025), the newt *Pleurodeles waltl* emerges as an ideal model to discover mechanisms of cellular plasticity in astrocyte-like glial cells, paving the way to the development of new cell type engineering approaches for brain repair.

## Acknowledgements

We thank the Columbia University ICM team for the excellent animal care, all members of the Tosches lab and Joanna Smeeton, Hynek Wichterle, Dritan Agalliu, and Bianca Dumitrascu for critical input throughout the project. We thank Humberto Ibarra Avila from the Zuckerman Institute Cellular Imaging Facility for support, Saket Choudhary for advice on bioinformatics, Nicholas J. Chua for help with scRNAseq data remapping, and Nastasia Nelson and Njeri Z. R. Sparman for preliminary HCR data. Computing resources were provided by Columbia University’s Shared Research Computing Facility (NIH grant 1G20RR030893-01 and NYSTAR contract C090171). This work was supported by a Columbia University RISE Seed Funding grant (MAT), the National Institute of Health (grant R35GM146973 to MAT), the Rita Allen Foundation (MAT), the Chan Zuckerberg Initiative (MAT), and the Pershing Square Foundation (MAT).

## Author contributions

A.O-G. and M.A.T. designed the study. A.O-G., M.K., D.A. and J.W. performed experiments. A.O-G., M.K. and M.A.T. analyzed the data. A.O-G., M.K. and M.A.T. wrote the manuscript with input from all authors. M.A.T. obtained funding and supervised the study.

## Materials and Methods

### Animals

*Pleurodeles waltl* larvae and adults were obtained from a breeding colony established at Columbia University. The animals were maintained in an aquatics facility at 20-25 ºC under a 12L:12D cycle. All experiments were conducted in accordance with the NIH guidelines and with the approval of the Columbia University Institutional Animal Care and Use Committee (IACUC protocols AC-AABI2617 and AC-AABF2564). Larvae were staged according to (Gallien & Durocher, 1957).

### Single cell dissociation and sequencing

To expand the dataset generated by (Woych et al., 2022), we dissociated the telencephalon from two adult animals. Single cell suspensions were obtained according to (Woych et al., 2022). All dissociated cells were multiplexed usingCell Multiplexing Oligos (10x Genomics 3’ CellPlex Multiplexing kit) prior to GEM formation. Sequencing libraries were prepared according to the 10x Genomics 3’ Next GEM 3’ v3.1 kit instructions, and sequenced on a NovaSeq X Plus platform with 150 bp paired-end reads.

### Pallial injury survival surgery

Adult and larval animals were anesthetized by immersion in 0.1% and 0.02% MS-222, respectively, buffered in a pH 7.4 tank water solution. Adult animals were placed in a custom 3D-printed holding chamber that fitted into a petri dish. Animals were covered with a sterile gauze previously soaked in anesthetic solution. Using a scalpel and fine forceps, a skin flap was cut above the brain and carefully detached from the skull; the surface of the skull was then scraped with the scalpel. A cotton-tip applicator was used to dry the surface of the skull and a dental drill was used to make a craniotomy above the left hemisphere, spanning the entire rostro-caudal length of the choroid plexus. The three meningeal membranes were then removed carefully with fine forceps and scissors. Once the surface of the brain was clear from membranes, a biopsy punch device (Stoelting 57391) 250 µm in diameter was used. To target the same pallial region in a reproducible way, we used the most anterior edge of the choroid plexus as a landmark, and placed the biopsy punch approximately 150 µm lateral to it. The biopsy punch was then carefully pushed down for about 200 µm from the surface of the brain, to remove a portion of the pallium spanning its entire thickness. For surgeries with an experimental endpoint of more than 10 days, a round glass coverslip 3 mm in diameter was used to cover the craniotomy. The glass coverslip was secured in place using dental cement. Once the cement dried, the skin flap was placed back on the skull, and fixed with a small amount of silicone adhesive (WPI KWIKSIL). Animals were allowed to recover in tanks with moisturized towels but no water for 24 hours, after which they were monitored and transitioned to a tank full of water until the experimental endpoint.

Injuries in larvae were carried out similarly. However, larvae were placed in 100 mm petri dishes with sterile gauze pads soaked in 0.02% MS-222 buffered in a pH 7.4 tank water solution for more grip. The brain was exposed by cutting the skull along the midline with a scalpel. Two horizontal incisions in the skull above the left hemisphere were made with fine scissors, going laterally from the midline incision. The first incision was at the anterior boundary of the choroid plexus and the second incision was made at the most posterior limit of the choroid plexus. The resulting piece of skull was then removed from the head of the animal. After removing the meninges and making the injury with the biopsy punch, the skin flap was placed back on the head of the larvae and air-dried for 5 minutes. The animal was returned to the tank full of water, where it recovered until the experimental endpoint.

### Electroporation

Glass capillary needles were pulled and then broken open using fine forceps (needle tip diameter ranging 5-15 um), backfilled with mineral oil, and then connected to a Nanoject III injection system (Drummond), and filled with plasmid solution. Right before injection, adult animals were anesthetized in 0.1% MS-222. The brain was accessed as described for the pallial injury surgery. The needle was then inserted in the right lateral telencephalic ventricle, and plasmid solution was pressure injected. A volume of 600 nL plasmid solution was pressure injected at 50 nL/s. For pCAG-mGFP electroporation, a plasmid DNA solution of 1.5 ug/uL was prepared. FastGreen was added to the plasmid prep at a 1:10 ratio to visualize the injected solution.

Immediately after injection, the head was placed between electroporation electrodes (round 5 mm diameter platinum plate electrodes attached to tweezers (Nepagene, CUY650P5). The electrodes were covered with 4% agarose and trimmed to allow a ∼2mm distance from the head to the electrode. The positive electrode was placed dorsally and electroporations were performed with a BTX electroporator (ECM830). Five 50 ms unidirectional pulses with a 1 s interval were applied at 55, as described in (Joven et al., 2018).

### EdU administration

Larvae and adult animals were anesthetized in 0.1% and 0.02% MS-222, respectively, buffered in a pH 7.4 tank water solution. Animals were placed on their back in a petri dish and a single EdU injection (2.5 or 5 mg/mL stock solution in sterile 0.9% saline) was administered intraperitoneally (50 mg/kg of total body weight), using an insulin syringe.

### Tissue processing for whole mount IHC and HCR

For non-survival surgeries, animals were anesthetized by immersion in MS-222 as described above (0.2% MS-222 used for adults and 0.02% for larvae). For larvae at stage 50 or older, animals were transcardially perfused with ice-cold molecular-grade phosphate-buffered saline (PBS) before brain dissection. Adult animals were perfused with PBS and then with ice-cold 4% paraformaldehyde (PFA) in PBS before brain dissection. After dissection, brains were fixed overnight in 4% PFA in PBS at 4 °C. Next, brains were dehydrated (20%, 40%, 60%, 80%, 100%, 100% MeOH in PBS, 45 min each at room temperature), placed in fresh 100% methanol, and stored at -20°C until further processing.

HCRs with and without EdU detection were performed as described in (Jaeger et al., 2025). Specifically, brains were first stained as whole-mount preparations, then embedded in 4% agarose in Tris-HCl (500 mM, pH 7.0), and finally sectioned to obtain 70 µm-thick sections for confocal imaging. EdU was detected using the Click-IT reaction kit (Invitrogen), according to the manufacturer instructions. If combining immunofluorescence with HCR, sections were incubated with primary antibody solution (Tris-HCl) for 1 - 3 nights at 4º C. Tissue sections were then washed 3 × 20 min in Tris-HCl at RT, incubated with secondary antibody solution (DAPI 1:1000 in Tris-HCl) overnight hrs at 4º C, and again washed for 3 × 20 min before mounting in Fluoromount-G® Mounting Medium (SouthernBiotech) or DAKO fluorescent mounting medium (Agilent). All images were acquired using a Zeiss LSM800 confocal microscope. HCR-3.0-style probe pairs against *Gja1, Sox4, Aqp4, Eomes, Nts, Slc17a7, Frmd8, and Satb1* were designed using insitu_probe_generator (Kuehn et al., 2022) and ordered from Integrated DNA Technologies, and 6 pmol of probe was added to the probe hybridization buffer (Molecular Instruments) for incubation. Antibodies used were the rabbit anti-Sox2 (1:500, Abcam ab97959), and the goat anti-rabbit IgG Alexa 594 as secondary (1:500, Invitrogen A-11037).

A subset of EdU-stained brains was imaged as whole-mounts with a light-sheet microscope. In this case, stained brains were dehydrated after staining in 50% MeOH in TrisHCl for 1 hr at RT, followed by two 1 hour 100% MeOH washes at RT. Samples were then incubated in 66% dichloromethane (DCM)/ 33% MeOH for 3 hrs at RT. Residual methanol was removed with 2 × 15 min washes in 100% DCM, and the tissue was allowed to clear overnight in dibenzyl ether (DBE). Images were acquired using a LaVision Ultramicroscope II light sheet microscope with a 4X DBE immersion objective and 2 µm pixel resolution, and visualized using ImarisViewer 9.8.0.

### 6-propyl-2-thiouracil (PTU) treatment

Stage 54 larvae were identified according to (Gallien & Durocher, 1957) and divided in two experimental groups: PTU treated (n = 22 animals) and control siblings (n = 19 animals). A stock solution of 40 L was prepared by dissolving PTU in tank water at a concentration of 50 mg/L at least once a month. This stock solution was used to replace PTU-treated water in the animal tanks every two days. Treated animals were kept in PTU-water from the onset of stage 54 (∼79 day-old animals) until euthanasia, for a total duration of 51 weeks. Metamorphosis occurred naturally in the control (untreated) siblings.

### Confocal microscopy and quantifications

Sections imaged with the Zeiss LSM800 confocal microscope were processed in Fiji. All cell counts reported in the paper were done manually using the Cell Counter Plugin in Fiji.

#### Proliferation and neurogenesis after pallial injury

To quantify the number of *Eomes+* and EdU*+* cells in the injured and contralateral uninjured hemispheres, 1 section at the site of the injury was imaged as tiled z-stacks (7 µm distance between imaging planes). All *Eomes*+ and EdU+ cells were counted across 3 central imaging planes in 100 µm-wide regions (measured along the ventricular surface) on the medial and on the lateral sides of the injury. The *Eomes+* and EdU*+* cells in the contralateral hemisphere were counted within a 200 µm-wide area. The data is presented as mean ± SD (for larval condition: n = 5 animals for 3dpi, 7dpi, 10dpi, 15dpi; n = 6 animals for 5dpi; n = 4 for 21dpi. For adult condition: n = 4 animals for 3dpi, 7dpi, 15dpi, 21dpi; n = 5 animals for 5dpi; n = 8 animals for 10dpi. For PTU experiment: n = 5 at 10dpi for each condition).

#### Adult neurogenesis

Brains were stained for DAPI, *Nts, Frmd8*, Sox2, and EdU. These stains were imaged with the following lasers: 405 nm (DAPI), 488 nm (EdU), 546 nm (*Frmd8* HCR), 594 nm (Sox2 antibody), and 647 nm (*Nts* HCR). For each brain, eight coronal sections of the anterior dorsal pallium spanning its rostro-caudal axis were imaged as tiled z-stacks, with a 7-µm distance between imaging planes. All EdU+ cells in the DP and LP were manually counted and scored for expression of *Nts, Frmd8*, and Sox2. The boundary between the DP and MP was established from anatomical landmarks (tissue curvature, cell density). The boundary between the LP and VP was established from anatomical landmarks and from the presence of *Nts* in the VP but not in LP. Since HCRs had to be imaged at high laser power, we observed signal bleedthrough from the Sox2 staining (antibody, nuclear signal) into the 546 nm channel (*Frmd8*, HCR, cytoplasmic signal). Our scRNAseq data and independent stainings show that these two genes are not coexpressed in the same cells in the pallium. Therefore, we used the following criteria to count cells: all the cells with a nuclear staining (Sox2) in the 594 nm channel were classified as EGCs if close to the ventricular surface and as GABAergic interneurons if parenchymal; all the cells with a cytoplasmic stain in the 546 nm channel that were not labeled also in the 594 nm channel were classified as DP-DL *Frmd8*+ neurons. Cell proportions were then calculated, and data presented as mean ± SD (n = 3 animals analyzed for each time point). For final visualization, we masked the Sox2 signal from the *Frmd8* signal with Fiji image subtraction.

#### Regeneration of neuron types

Brains were stained for DAPI, *Nts, Satb1*, Sox2, and EdU. These stains were imaged with the following lasers: 405 nm (DAPI), 488 nm (EdU), 546 nm (*Satb1* HCR), 594 nm (Sox2 antibody), and 647 nm (*Nts* HCR). For each brain, 3 vibratome sections spanning the injury site (70 µm-thick sections, for a total of ∼210 µm) were imaged as tiled z-stacks (7 µm interval; 10 imaging planes on average). All EdU+ cells in the DP and LP were manually counted in three consecutive imaging planes from each z-stack and scored for expression of *Nts, Satb1* and *Sox2*. The same strategy described above was used to distinguish *Satb1* signal (HCR, 546 nm) and *Sox2* signal (antibody, 594 nm). Cell proportions were then calculated, and data presented as mean ± SD (n = 5 animals analyzed for each condition).

To quantify the number of *Satb1*+ neurons in the regenerated pallium and the contralateral pallium, we used the same data z-stacks and same sections described above, counting all *Satb1*+ cells regardless of EdU incorporation. All *Satb1*+ cells co-labeled with Sox2 were not counted as LP *Satb1*+ neurons, in accordance with our scRNAseq data showing that *Satb1* and Sox2 are coexpressed in GABAergic interneurons, and not in LP cells. Cell ratios were then calculated by dividing the total number of *Satb1*+ cells in the injured hemisphere by the total number of *Satb1+* cells in the contralateral uninjured hemisphere. The data is presented as mean ± SD (n = 3 animals analyzed for each condition).

To quantify the ratio of DP-SL neurons across the injured and the contralateral hemisphere, the middle imaging plane from the same z-stacks used to count EdU+ cells (1 imaging plane per section; a total of 3 sections per animal) were used. All *Nts*+ cells in the DP were counted regardless of EdU incorporation. All *Nts*+ cells co-labeled with Sox2 were not counted as DP-SL, in accordance with our scRNAseq data showing that *Nts* and Sox2 are coexpressed in GABAergic interneurons, and not in DP-SL cells. Cell ratios were then calculated by dividing the total number of *Nts*+ cells in the injured hemisphere by the total number of *Nts*+ cells in the contralateral uninjured hemisphere. The total number of DP-DL neurons in two hemispheres was obtained by exclusion, by counting all the cells (DAPI-stained nuclei) in the pallium neuronal layer that were not labeled by the DP-SL marker (*Nts*), the LP marker (*Satb1*) and the neural progenitor and GABAergic interneuron marker (Sox2); the ratio between hemispheres was calculated as described above. The data is presented as mean ± SD (n = 5 animals analyzed for each condition). Similar to the adult neurogenesis images (**Figure6**), we masked the Sox2 signal from the *Satb1* signal with Fiji image subtraction for final visualization (see **Supplementary Figure 5C**).

### Statistical analysis

Statistical analyses on EdU and *Eomes* cell counts after pallial injury were performed using R (version 4.4.0) with the emmeans (1.11.2-8) and dplyr (1.1.4) packages. To compare changes in EdU+ and Eomes+ cell populations between larvae and adults, we performed two-way ANOVAs with condition (larvae vs adult) and timepoint as factors in the injured hemisphere. To evaluate injury response relative to baseline, we performed separate two-way ANOVAs for each condition (larvae and adults) with timepoint and hemisphere as factors. Lastly, to assess regional differences in the adult regenerative response, we performed a two-way ANOVA with timepoint and region (lateral vs medial) as factors. Post-hoc pairwise comparisons were conducted using estimated marginal means (EMMs) with Bonferroni correction for multiple comparisons. Statistical significance was set at α = 0.05. All p-values reported in the text are adjusted for multiple comparisons using the Bonferroni method.

For PTU-treatment, EdU+ and *Eomes*+ cell counts were compared between PTU-treated and control siblings separately for injured and contralateral regions using Welch’s two-sample t-tests, which do not assume equal variances between groups. Normality of data distribution was assessed using Shapiro-Wilk tests. Given that comparisons in injured and contralateral regions represent distinct biological contexts, and EdU+ and *Eomes*+ represent different cell populations, p-values are reported without correction for multiple comparisons. Statistical significance was set at α = 0.05.

The percentages and ratios of DP-SL and DP-SL neurons were compared between larval (n=5) and adult (n=5) animals using Welch’s two-sample t-tests, which do not assume equal variances between groups. Normality of data distribution within each group was assessed using the Shapiro-Wilk test, confirming that parametric testing was appropriate for both measures. The ratio of LP-*Satb1*+ neurons were compared between larval (n=3) and (n=3) animals using Welch’s two-sample t-tests. All analyses were performed in R (version 4.0.0). Given that percentage and ratio measures test distinct biological hypotheses (overall abundance vs. proportional composition), p-values are reported without correction for multiple comparisons. Statistical significance was set at α = 0.05.

### Analysis of single-cell RNA-seq data

#### Read alignment

Data used in this study was previously collected as described in (Woych et al., 2022) and (Deryckere et al., 2025); new scRNAseq data were added for the adult brain datasets. All reads, including from previous data, were aligned to a de-novo genome assembly of *Pleurodeles waltl* (aPleWal1.hap1.20221129) (Brown et al., 2025) using cellranger 8.0.1 and 9.0. The reference was supplemented with eGFP sequences for aligning the stage 55a/TrackerSeq libraries published in (Deryckere et al., 2025). An index was generated using the cellranger mkref command, using the fasta and GTF files provided in NCBI. The majority of developmental libraries were multiplexed using the 10x Genomics 3’ CellPlex Multiplexing kit, except for samples D1 (St. 41 dissociation) and D2 (St. 50 dissociation). For the adult libraries, only E7 and T1 were multiplexed using 10x Genomics 3’ CellPlex Multiplexing kit. For the libraries that required demultiplexing, we used cellranger multi command and multiplex tag reads were assigned. For the others, cellranger count command was used, using the transcriptome reference generated by cellranger and setting create-bam to true.

#### Data QC and filtering

Filtered count matrices obtained after alignment with cellranger were imported into Scanpy (1.11.4). Individual libraries were imported and filtered by keeping cells with ⋜15% of mitochondrial counts, >500 UMIs, >500 genes, and a number of genes < 3 SD above the mean. A single AnnData object was created from individual libraries and saved with Seurat compatibility.

Datasets were then read into Seurat 5.3.0 and filtered more extensively. We first removed all CMO tags that were carried as features in the count matrices. Next, after Louvain clustering, low-quality cells were removed using a Support Vector Machine (SVM) classifier (R package e1071) as described in (Tosches et al., 2018). Low-quality clusters (number of genes per cell below average of the dataset, percentage of mitochondrial genes above average of dataset, and absence of cluster-specific marker genes) were first identified, and 10% of cells from these clusters, together with 10% of cells from the rest of the dataset were used to train the classifier. Since part of this dataset was previously filtered as described in (Deryckere et al., 2025; Woych et al., 2022), we included those cell names in the top-quality cells set. The trained SVM classifier then identified low-quality cells across the dataset, which we removed to obtain a final Seurat object of 132,548 cells in the developmental dataset and 67,541 cells in the adult dataset.

#### Pre-processing and label transfer

Both datasets were clustered and analyzed using the R package Seurat 5.3.0 (Hao et al., 2024). CellCycleScoring was applied on both datasets, after which the raw counts were normalized using SCTransform (Choudhary & Satija, 2022; Hafemeister & Satija, 2019), while regressing the data for the number of UMIs per cell, developmental stage, percent mitochondrial genes, library source, multiplexed condition, G2M and S scores for the developmental object; and number of UMIs per cell, stage, library source, multiplexed condition, housing condition, G2M and S scores for the adult object. For both datasets, the top 3000 variable genes were used and a principal component analysis (PCA) was performed, after which the first 180 principal components were used for clustering and UMAP embedding the adult dataset, while 125 principal components were used for the developmental dataset. The default Louvain clustering method was used, with a resolution of 2 in both datasets. This pipeline allowed us to recover 97.23% of all the previously published cells in (Woych et al., 2022), as well as 91.14% of the previously published cells in (Deryckere et al., 2025).

To annotate the adult object, Seurat’s label transfer was performed using the previously published dataset with updated annotations made in (Deryckere et al., 2023, 2025). The Label Transfer algorithm identifies transfer anchors between two data sets, allowing comparison of cell type identities between the two. For this transfer, the function FindTransferAnchors was run using dims = 50, and k.anchor = 10 and the function TransferData was run using k.weight = 20. For the developmental object, we took the same approach that was used in (Deryckere et al., 2025), using iterative label transfer to maximize the matching of cell types. We used the same reference as before for querying the stage 55 cells in the developmental object. Subsequently, the annotated stage 55 dataset was used as a reference for identifying cell identity in the stage 53 object, and so on. For all transfers within the developmental object, the function FindTransferAnchors was run using dims = 20, and k.anchor = 5 and the function TransferData was run using k.weight = 20.

#### Heterogeneity of pallial progenitors

For subsetting the neurogenic trajectories in larvae and adult, telencephalic *Foxg1*-expressing cells were identified, and then we removed cells expressing *Otp* and *Sim1*, corresponding to cells from the hypothalamus and the pre-optic area (POA) (Deryckere et al., 2023). Immature neurons and neural stem cells and intermediate progenitors were identified on the basis of *Sox9, Slc1a3, Gfap, Sox2*, and *Fabp7* expression (for NSCs) and high expression of *Sox4, Mex3a* and low expression of *Snap25* and *Syt1* (for immature neurons). The final dataset contained 33,072 cells, of which 30,750 cells were from larvae and 2,322 cells were from adults. For subsetting pallial progenitors from both larval and adult progenitors, we used an iterative approach. Subpallial cells were removed on the basis of *Gad2, Fzd5, Dlx2, Vax1, Gsx2* and *Nkx2*.*1* expression. The remaining cells were corroborated to be pallial cells by expression of *Emx1* and *Pax6*, and in addition, *Sox6*+ cells were included; these cells correspond to the ventral pallium (*Emx1* negative (Brox et al., 2004)). Lastly, from this pallial dataset, we subsetted the progenitors based on high *Slc1a3, Gfap, Sox2* and *Fabp7* expression and absence of *Neurog2, Tubb3* and *Snap25* (*i*.*e*. not yet committed to a neuronal fate). The same procedure was applied to both adult and developmental datasets. After validating the pallial nature of the subsetted progenitors, the raw count matrices from pre- and adult pallial progenitors subset were merged into a single Seurat object. This final object contained 6,959 cells, including for 2,214 adult progenitors and 4,745 larval progenitors. We next performed a principal component analysis (PCA), after regressing the data for number of UMIs per cell, number of genes per cell, stage, library source, multiplexed condition, housing condition, life cycle stage, G2M and S scores. The top 3000 variable genes were used as well as the first 75 principal components for clustering and UMAP embedding. Finally the default Louvain clustering method was used, with a resolution of 1.

We then annotated the pallial progenitors depending on the pallial field to which they belonged to, following the same approach as (Deryckere et al., 2025), using previously generated custom functions. For that, we fit a principal curve that represents an ordered ‘spatial trajectory’ for each cell based on their expression profile (Hastie & Stuetzle, 1989), using the principal_curve function in the princurve R Package with options smoother = ‘lowess’, f = ⅓ and stretch=2 (Cannoodt & Bengtsson, 2019). Consistently, we defined a ‘mediolateral score’ as the length of the arc from the beginning of the curve to the point where the cell projects onto the curve. Since there is no directionality in PCs, we defined an explicit starting point based on the expression of *Wnt8b*, such that its expression is negatively correlated with the principal curve score. We validated the ‘mediolateral score’ calculated from the principal curve by assessing the expression of genes along the principal curve that were previously validated to be spatially organized such as *Sfrp1, Wnt8b, Slit2, Dmrt3, Meis2* and *Sp8*. To identify genes that vary across the mediolateral axis, we used the custom function GetVaryingGenes2 (see (Deryckere et al., 2025)) that uses the fitGAM function from the tradeSeq package using default parameters. To limit the analysis to highly informative genes, we ran the function to limit only to those genes that were found to be high variable genes from the SCTranform function. To assign explicit spatial labels to the progenitors, we performed a hierarchical clustering using the genes that are differentially expressed across the mediolateral trajectory. We used the hclust() function with method=“ward.D2” to perform hierarchical clustering in R; the tree was cut at h=275 to obtain 11 clusters which were annotated based on gene expression of spatially-restricted markers (*Wnt8b, Slit2, Fezf2, Lhx9, Meis2, Sfrp1*), resulting in 5 groups of pallial progenitors: MP, DP, LP, VP, and AMY.

To characterize the activation state of the progenitors across brain regions, we used a scoring strategy similar to the mediolateral score. We fit a principal curve onto principal components 1 and 2, since the gene loading for these two PCs strongly suggested a gradient from quiescence to active proliferation. We then defined an ‘activation score’ as the length of the arc from the beginning of the curve to the point where the cell projects onto the curve. Genes differentially expressed along the activation trajectory were identified with the GetVaryingGenes2 function, as described above. We then performed hierarchical clustering (hclust function, method=“ward.D2”), with a tree cutoff at h=1500 to obtain 3 cell state clusters, annotated on the basis of gene expression of cell cycle related genes: deep quiescence (dQ), shallow quiescence (sQ), and active (A).

#### Orthologous gene alignment

Only one-to-one orthologous genes were used for cross-species comparisons. EggNOG orthology assignments for newt (*Pleurodeles waltl*) and mouse (Mus musculus) were taken from (Woych et al., 2022).

#### Gene module scores in cross-species comparison

We analysed and compared the expression of cell-type specific gene modules to assess molecular similarity across cell types and species. To identify differentially-expressed genes (DEGs) in the newt dataset, we applied the Seurat (v5.3.0) FindMarkers function to perform pairwise comparisons of cell state clusters. Default settings (Wilcox) were chosen for the test.use parameter in the FindMarkers function. To obtain the top DEGs, we then filtered the DEG list to keep only genes expressed in at least 10% of the cells from at least one cell state cluster. Genes were classified as significantly enriched if they exhibited an adjusted p-value <0.05 and an absolute average log2 fold chance > 1.5. For each cell cluster, we took the top 100 genes DEGs in each comparison. After merging and filtering out duplicate genes, we obtained a final list of 136 marker genes for the active cluster, 165 genes for the shallow quiescence cluster and 144 genes for the deep quiescence cluster. Finally, these lists were filtered to keep only genes with a one-to-one ortholog in mouse. We finally used the AddModuleScore function in Seurat to examine the expression of these gene sets in mouse V-SVZ cell types from (Cebrian-Silla et al., 2025). A positive gene module score indicates that the gene set is highly expressed in a particular cell compared to the average expression across all cells. We visualized the results with UMAP plots as well as the distribution of the gene module score with Violin Plots.

For the analysis of mouse gene modules and of their expression in newt cells, we followed the same approach as described above but in opposite order. We computed DEGs on mouse astrocytes vs active B cells, astrocytes vs quiescent B cells, and active B cells vs quiescent B cells. Using the FindMarkers function with default (Wilcox) parameters, we filtered the list to keep only genes expressed in at least 10% of the cells and considered significant if they exhibited an adjusted p-value <0.05 and an absolute average log2 fold change > 1.5. Then we took only the top 100 DEGs for each cell cluster and comparison. When comparing quiescent B cells to parenchymal astrocytes we noticed that all of the genes upregulated in quiescent B cells were related to ribosomal genes and translation. Since most of the DEGs in quiescent B cells compared to astrocytes were ribosomal genes, we excluded this list from the computation. The resulting gene sets contained 97 unique genes for active B cells, 92 unique genes for astrocytes and 95 unique genes for quiescent B cells. We then filtered these lists further to keep only genes with a one-to-one ortholog in newt, and finally computed gene module scores using Seurat’s AddModuleScore function in the newt dataset.

## References

Alibardi, L. (2023). Regeneration among animals: An evolutionary hypothesis related to aquatic versus terrestrial environment. Developmental Biology, 501, 74–80. 10.1016/j.ydbio.2023.06.013

Alunni, A., & Bally-Cuif, L. (2016). A comparative view of regenerative neurogenesis in vertebrates. Development, 143(5), 741–753. 10.1242/dev.122796

Amamoto, R., Huerta, V. G. L., Takahashi, E., Dai, G., Grant, A. K., Fu, Z., & Arlotta, P. (2016). Adult axolotls can regenerate original neuronal diversity in response to brain injury. eLife, 5, e13998. 10.7554/eLife.13998

Arsanto, J.-P., Komorowski, T. E., Dupin, F., Caubit, X., Diano, M., Géraudie, J., Carlson, B. M., & Thouveny, Y. (1992). Formation of the peripheral nervous system during tail regeneration in urodele amphibians: Ultrastructural and immunohistochemical studies of the origin of the cells. Journal of Experimental Zoology, 264(3), 273–292. 10.1002/jez.1402640307

Ayana, R., Zandecki, C., Van houcke, J., Mariën, V., Seuntjens, E., & Arckens, L. (2024). Single-cell sequencing unveils the impact of aging on the progenitor cell diversity in the telencephalon of the female killifish N. furzeri. Aging Cell, 23(10), e14251. 10.1111/acel.14251

Becker, C. G., & Becker, T. (2015). Neuronal Regeneration from Ependymo-Radial Glial Cells: Cook, Little Pot, Cook! Developmental Cell, 32(4), 516–527. 10.1016/j.devcel.2015.01.001

Berg, D. A., Kirkham, M., Beljajeva, A., Knapp, D., Habermann, B., Ryge, J., Tanaka, E. M., & Simon, A. (2011). Efficient regeneration by activation of neurogenesis in homeostatically quiescent regions of the adult vertebrate brain. Development, 138(1), 180. 10.1242/dev.061754

Blackshaw, S. (2022). Why Has the Ability to Regenerate Following CNS Injury Been Repeatedly Lost Over the Course of Evolution? Frontiers in Neuroscience, 16. 10.3389/fnins.2022.831062

Brown, T., Mishra, K., Elewa, A., Iarovenko, S., Subramanian, E., Araus, A. J., Petzold, A., Fromm, B., Friedländer, M. R., Rikk, L., Suzuki, M., Suzuki, K. T., Hayashi, T., Toyoda, A., Oliveira, C. R., Osipova, E., Leigh, N. D., Yun, M. H., & Simon, A. (2025). Chromosome-scale genome assembly reveals how repeat elements shape non-coding RNA landscapes active during newt limb regeneration. Cell Genomics, 5(2).

Brox, A., Puelles, L., Ferreiro, B., & Medina, L. (2004). Expression of the genes Emx1, Tbr1, and Eomes (Tbr2) in the telencephalon of Xenopus laevis confirms the existence of a ventral pallial division in all tetrapods. Journal of Comparative Neurology, 474(4), 562–577. 10.1002/cne.20152

Cannoodt, R., & Bengtsson, H. (2019). rcannood/princurve: Princurve 2.1.4 [Computer software]. Zenodo. 10.5281/zenodo.3351282

Cebrian-Silla, A., Nascimento, M. A., Mancia, W., Gonzalez-Granero, S., Romero-Rodriguez, R., Obernier, K., Steffen, D. M., Lim, D. A., Garcia-Verdugo, J. M., & Alvarez-Buylla, A. (2025). Neural stem cell relay from B1 to B2 cells in the adult mouse ventricular-subventricular zone. Cell Reports, 44(3). 10.1016/j.celrep.2025.115264

Chen, J., Poskanzer, K. E., Freeman, M. R., & Monk, K. R. (2020). Live-imaging of astrocyte morphogenesis and function in zebrafish neural circuits. Nature Neuroscience, 23(10), 1297–1306. 10.1038/s41593-020-0703-x

Choudhary, S., & Satija, R. (2022). Comparison and evaluation of statistical error models for scRNA-seq. Genome Biology, 23(1), 27. 10.1186/s13059-021-02584-9

Clinton, B. K., Cunningham, C. L., Kriegstein, A. R., Noctor, S. C., & Martínez-Cerdeño, V. (2014). Radial glia in the proliferative ventricular zone of the embryonic and adult turtle, Trachemys scripta elegans. Neurogenesis, 1(1), e970905. 10.4161/23262125.2014.970905

Cosacak, M. I., Bhattarai, P., Reinhardt, S., Petzold, A., Dahl, A., Zhang, Y., & Kizil, C. (2019). Single-Cell Transcriptomics Analyses of Neural Stem Cell Heterogeneity and Contextual Plasticity in a Zebrafish Brain Model of Amyloid Toxicity. Cell Reports, 27(4), 1307–1318.e3. 10.1016/j.celrep.2019.03.090

Degitz, S. J., Holcombe, G. W., Flynn, K. M., Kosian, P. A., Korte, J. J., & Tietge, J. E. (2005). Progress towards Development of an Amphibian-Based Thyroid Screening Assay Using Xenopus laevis. Organismal and Thyroidal Responses to the Model Compounds 6-Propylthiouracil, Methimazole, and Thyroxine. Toxicological Sciences, 87(2), 353–364. 10.1093/toxsci/kfi246

Deryckere, A., Choudhary, S., Lynch, C., Limperis, L. K. V. P., Affatato, P., Woych, J., Gumnit, E., Ortega Gurrola, A., Satija, R., Mayer, C., & Tosches, M. A. (2025). A conserved logic for the development of cortical layering in tetrapods (p. 2025.10.01.679862). bioRxiv. 10.1101/2025.10.01.679862

Deryckere, A., Woych, J., Jaeger, E. C. B., & Tosches, M. A. (2023). Molecular Diversity of Neuron Types in the Salamander Amygdala and Implications for Amygdalar Evolution. Brain Behavior and Evolution, 98(2), 61–75. 10.1159/000527899

Echeverri, K., & Tanaka, E. M. (2002). Ectoderm to Mesoderm Lineage Switching During Axolotl Tail Regeneration. Science, 298(5600), 1993–1996. 10.1126/science.1077804

Elewa, A., Wang, H., Talavera-López, C., Joven, A., Brito, G., Kumar, A., Hameed, L. S., Penrad-Mobayed, M., Yao, Z., Zamani, N., Abbas, Y., Abdullayev, I., Sandberg, R., Grabherr, M., Andersson, B., & Simon, A. (2017). Reading and editing the Pleurodeles waltl genome reveals novel features of tetrapod regeneration. Nature Communications, 8(1), 2286. 10.1038/s41467-017-01964-9

Filoni, S., Bernardini, S., & Cannata, S. M. (1995). Differences in the decrease in regenerative capacity of various brain regions of Xenopus laevis are related to differences in the undifferentiated cell populations. Journal Fur Hirnforschung, 36(4), 523–529.

Frangioni, G., Atzori, A., Balzi, M., Fuzzi, G., Ghinassi, A., Pescosolido, N., Bianchi, S., & Borgioli, G. (2006). Thyroid and hypoxic stress in the newt Triturus carnifex. Journal of Experimental Zoology Part A: Comparative Experimental Biology, 305A(3), 225–232. 10.1002/jez.a.268

Fu, S., Zeng, Y.-Y., Peng, C., Wang, L., Feng, Y., Wang, K., Liu, Y., & Fei, J.-F. (2026). Ependymoglial cells are critical for cortex regeneration in axolotls. Nature Communications, 17(1), 1827. 10.1038/s41467-026-68538-6

Gallien, L., & Durocher, M. (1957). Table chronologique du développement chez Pleurodeles waltlII Michah. 2, 1–19.

Goldshmit, Y., Sztal, T. E., Jusuf, P. R., Hall, T. E., Nguyen-Chi, M., & Currie, P. D. (2012). Fgf-Dependent Glial Cell Bridges Facilitate Spinal Cord Regeneration in Zebrafish. Journal of Neuroscience, 32(22), 7477–7492. 10.1523/JNEUROSCI.0758-12.2012

Gumnit, E., Gattoni, G., Woych, J., Deryckere, A., Bhattarai, P., Gillis, J. A., Kizil, C., & Tosches, M. A. (2026). Evolutionary origins and transcriptomic innovations of vertebrate Cajal-Retzius cells. Current Biology, 36(9), 2397–2412.e7. 10.1016/j.cub.2026.03.082

Hafemeister, C., & Satija, R. (2019). Normalization and variance stabilization of single-cell RNA-seq data using regularized negative binomial regression. Genome Biology, 20(1), 296. 10.1186/s13059-019-1874-1

Hao, Y., Stuart, T., Kowalski, M. H., Choudhary, S., Hoffman, P., Hartman, A., Srivastava, A., Molla, G., Madad, S., Fernandez-Granda, C., & Satija, R. (2024). Dictionary learning for integrative, multimodal and scalable single-cell analysis. Nature Biotechnology, 42(2), 293–304. 10.1038/s41587-023-01767-y

Hastie, T., & Stuetzle, W. (1989). Principal Curves. Journal of the American Statistical Association, 84(406), 502–516. 10.1080/01621459.1989.10478797

Hayashi, T., Yokotani, N., Tane, S., Matsumoto, A., Myouga, A., Okamoto, M., & Takeuchi, T. (2013). Molecular genetic system for regenerative studies using newts. Development, Growth & Differentiation, 55(2), 229–236. 10.1111/dgd.12019

Hirose, K., Payumo, A. Y., Cutie, S., Hoang, A., Zhang, H., Guyot, R., Lunn, D., Bigley, R. B., Yu, H., Wang, J., Smith, M., Gillett, E., Muroy, S. E., Schmid, T., Wilson, E., Field, K. A., Reeder, D. M., Maden, M., Yartsev, M. M., … Huang, G. N. (2019). Evidence for hormonal control of heart regenerative capacity during endothermy acquisition. Science, 364(6436), 184–188. 10.1126/science.aar2038

Hui, S. P., Dutta, A., & Ghosh, S. (2010). Cellular response after crush injury in adult zebrafish spinal cord. Developmental Dynamics, 239(11), 2962–2979. 10.1002/dvdy.22438

Jaeger, E. C. B., Vijatovic, D., Deryckere, A., Zorin, N., Nguyen, A. L., Ivanian, G., Woych, J., Arnold, R. C., Gurrola, A. O., Shvartsman, A., Barbieri, F., Toma, F. A., Cline, H. T., Shay, T. F., Kelley, D. B., Yamaguchi, A., Shein-Idelson, M., Tosches, M. A., & Sweeney, L. B. (2025). Adeno-associated viral tools to trace neural development and connectivity across amphibians. Developmental Cell, 60(5), 794–812.e6. 10.1016/j.devcel.2024.10.025

Joven, A., Elewa, A., & Simon, A. (2019). Model systems for regeneration: Salamanders. Development, 146(14), dev167700. 10.1242/dev.167700

Joven, A., Wang, H., Pinheiro, T., Hameed, L. S., Belnoue, L., & Simon, A. (2018). Cellular basis of brain maturation and acquisition of complex behaviors in salamanders. Development, 145(1), dev160051. 10.1242/dev.160051

Kaslin, J., Kroehne, V., Benato, F., Argenton, F., & Brand, M. (2013). Development and specification of cerebellar stem and progenitor cells in zebrafish: From embryo to adult. Neural Development, 8(1), 9. 10.1186/1749-8104-8-9

Kirkham, M., Hameed, L. S., Berg, D. A., Wang, H., & Simon, A. (2014). Progenitor Cell Dynamics in the Newt Telencephalon during Homeostasis and Neuronal Regeneration. Stem Cell Reports, 2(4), 507–519. 10.1016/j.stemcr.2014.01.018

Kroehne, V., Freudenreich, D., Hans, S., Kaslin, J., & Brand, M. (2011). Regeneration of the adult zebrafish brain from neurogenic radial glia-type progenitors. Development, 138(22), 4831–4841. 10.1242/dev.072587

Kuehn, E., Clausen, D. S., Null, R. W., Metzger, B. M., Willis, A. D., & Özpolat, B. D. (2022). Segment number threshold determines juvenile onset of germline cluster expansion in Platynereis dumerilii. Journal of Experimental Zoology Part B: Molecular and Developmental Evolution, 338(4), 225–240. 10.1002/jez.b.23100

Lange, C., Rost, F., Machate, A., Reinhardt, S., Lesche, M., Weber, A., Kuscha, V., Dahl, A., Rulands, S., & Brand, M. (2020). Single cell sequencing of radial glia progeny reveals the diversity of newborn neurons in the adult zebrafish brain. Development, 147(1), dev185595. 10.1242/dev.185595

Laywell, E. D., Rakic, P., Kukekov, V. G., Holland, E. C., & Steindler, D. A. (2000). Identification of a multipotent astrocytic stem cell in the immature and adult mouse brain. Proceedings of the National Academy of Sciences, 97(25), 13883–13888. 10.1073/pnas.250471697

Lust, K., Maynard, A., Gomes, T., Fleck, J. S., Camp, J. G., Tanaka, E. M., & Treutlein, B. (2022). Single-cell analyses of axolotl telencephalon organization, neurogenesis, and regeneration. Science, 377(6610), eabp9262. 10.1126/science.abp9262

Lust, K., & Tanaka, E. M. (2019). A Comparative Perspective on Brain Regeneration in Amphibians and Teleost Fish. Developmental Neurobiology, 79(5), 424–436. 10.1002/dneu.22665

Lyu, P., Iribarne, M., Serjanov, D., Zhai, Y., Hoang, T., Campbell, L. J., Boyd, P., Palazzo, I., Nagashima, M., Silva, N. J., Hitchcock, P. F., Qian, J., Hyde, D. R., & Blackshaw, S. (2023). Common and divergent gene regulatory networks control injury-induced and developmental neurogenesis in zebrafish retina. Nature Communications, 14(1), 8477. 10.1038/s41467-023-44142-w

Maden, M., Manwell, L. A., & Ormerod, B. K. (2013). Proliferation zones in the axolotl brain and regeneration of the telencephalon. Neural Development, 8(1), 1. 10.1186/1749-8104-8-1

Marqués-Torrejón, M. Á., Williams, C. A. C., Southgate, B., Alfazema, N., Clements, M. P., Garcia-Diaz, C., Blin, C., Arranz-Emparan, N., Fraser, J., Gammoh, N., Parrinello, S., & Pollard, S. M. (2021). LRIG1 is a gatekeeper to exit from quiescence in adult neural stem cells. Nature Communications, 12(1), 2594. 10.1038/s41467-021-22813-w

Matheson, A. M. M., Chua, N. J., & Tosches, M. A. (2025). Iberian ribbed newts. Current Biology, 35(2), R49–R51. 10.1016/j.cub.2024.11.062

Mchedlishvili, L., Mazurov, V., Grassme, K. S., Goehler, K., Robl, B., Tazaki, A., Roensch, K., Duemmler, A., & Tanaka, E. M. (2012). Reconstitution of the central and peripheral nervous system during salamander tail regeneration. Proceedings of the National Academy of Sciences, 109(34), E2258–E2266. 10.1073/pnas.1116738109

Monaghan, J. R., Stier, A. C., Michonneau, F., Smith, M. D., Pasch, B., Maden, M., & Seifert, A. W. (2014). Experimentally induced metamorphosis in axolotls reduces regenerative rate and fidelity. Regeneration, 1(1), 2–14. 10.1002/reg2.8

Morizet, D., Foucher, I., Alunni, A., & Bally-Cuif, L. (2024). Reconstruction of macroglia and adult neurogenesis evolution through cross-species single-cell transcriptomic analyses. Nature Communications, 15(1), 3306. 10.1038/s41467-024-47484-1

Morizet, D., Foucher, I., Mignerey, I., Alunni, A., & Bally-Cuif, L. (2025). Notch signaling blockade links transcriptome heterogeneity in quiescent neural stem cells with reactivation routes and potential. Science Advances, 11(35), eadu3189. 10.1126/sciadv.adu3189

Otsuki, L., & Brand, A. H. (2018). Cell cycle heterogeneity directs the timing of neural stem cell activation from quiescence. Science, 360(6384), 99–102. 10.1126/science.aan8795

Otsuki, L., & Tanaka, E. M. (2022). Positional Memory in Vertebrate Regeneration: A Century’s Insights from the Salamander Limb. Cold Spring Harbor Perspectives in Biology, 14(6), a040899. 10.1101/cshperspect.a040899

Parish, C. L., Beljajeva, A., Arenas, E., & Simon, A. (2007). Midbrain dopaminergic neurogenesis and behavioural recovery in a salamander lesion-induced regeneration model. Development, 134(15), 2881–2887. 10.1242/dev.002329

Rodrigo Albors, A., Tazaki, A., Rost, F., Nowoshilow, S., Chara, O., & Tanaka, E. M. (2015). Planar cell polarity-mediated induction of neural stem cell expansion during axolotl spinal cord regeneration. eLife, 4, e10230. 10.7554/eLife.10230

Smirnov, S. V., Merkulova, K. M., & Vassilieva, A. B. (2020). Skull development in the Iberian newt, Pleurodeles waltl (Salamandridae: Caudata: Amphibia): timing, sequence, variations, and thyroid hormone mediation of bone appearance. Journal of Anatomy, 237(3), 543–555. 10.1111/joa.13210

Sperry, D. G., & Grobstein, P. (1985). Regulation of neuron numbers in Xenopus laevis: Effects of hormonal manipulation altering size at metamorphosis. Journal of Comparative Neurology, 232(3), 287–298. 10.1002/cne.902320302

Tanaka, E. M., & Ferretti, P. (2009). Considering the evolution of regeneration in the central nervous system. Nature Reviews Neuroscience, 10(10), 713–723. 10.1038/nrn2707

Tiwari, N., Pataskar, A., Péron, S., Thakurela, S., Sahu, S. K., Figueres-Oñate, M., Marichal, N., López-Mascaraque, L., Tiwari, V. K., & Berninger, B. (2018). Stage-Specific Transcription Factors Drive Astrogliogenesis by Remodeling Gene Regulatory Landscapes. Cell Stem Cell, 23(4), 557–571.e8. 10.1016/j.stem.2018.09.008

Tosches, M. A., Yamawaki, T. M., Naumann, R. K., Jacobi, A. A., Tushev, G., & Laurent, G. (2018). Evolution of pallium, hippocampus, and cortical cell types revealed by single-cell transcriptomics in reptiles. Science, 360(6391), 881–888. 10.1126/science.aar4237

Trujillo-Cenóz, O., Marichal, N., Rehermann, M. I., & Russo, R. E. (2014). The inner lining of the reptilian brain: A heterogeneous cellular mosaic. Glia, 62(2), 300–316. 10.1002/glia.22607

Tsissios, G., Theodoroudis-Rapp, G., Chen, W., Sallese, A., Smucker, B., Ernst, L., Chen, J., Xu, Y., Ratvasky, S., Wang, H., & Del Rio-Tsonis, K. (2023). Characterizing the lens regeneration process in Pleurodeles waltl. Differentiation, Ocular Development: A View from the Front to the Back of the Eye, 132, 15–23. 10.1016/j.diff.2023.02.003

Urata, Y., Yamashita, W., Inoue, T., & Agata, K. (2018). Spatio-temporal neural stem cell behavior leads to both perfect and imperfect structural brain regeneration in adult newts. Biology Open, 7(6), bio033142. 10.1242/bio.033142

Urbán, N., Blomfield, I. M., & Guillemot, F. (2019). Quiescence of Adult Mammalian Neural Stem Cells: A Highly Regulated Rest. Neuron, 104(5), 834–848. 10.1016/j.neuron.2019.09.026

Urbán, N., & Cheung, T. H. (2021). Stem cell quiescence: The challenging path to activation. Development, 148(3), dev165084. 10.1242/dev.165084

Valcárcel-Hernández, V., Mayerl, S., Guadaño-Ferraz, A., & Remaud, S. (2024). Thyroid hormone action in adult neurogliogenic niches: The known and unknown. Frontiers in Endocrinology, 15. 10.3389/fendo.2024.1347802

Velloso, F. J., Shankar, S., Parpura, V., Rakic, P., & Levison, S. W. (2022). Neural Stem Cells in Adult Mammals are not Astrocytes. ASN Neuro, 14(1), 17590914221134739. 10.1177/17590914221134739

Wei, X., Fu, S., Li, H., Liu, Y., Wang, S., Feng, W., Yang, Y., Liu, X., Zeng, Y.-Y., Cheng, M., Lai, Y., Qiu, X., Wu, L., Zhang, N., Jiang, Y., Xu, J., Su, X., Peng, C., Han, L., … Gu, Y. (2022). Single-cell Stereo-seq reveals induced progenitor cells involved in axolotl brain regeneration. Science, 377(6610), eabp9444. 10.1126/science.abp9444

Woych, J., Ortega Gurrola, A., Deryckere, A., Jaeger, E. C. B., Gumnit, E., Merello, G., Gu, J., Joven Araus, A., Leigh, N. D., Yun, M., Simon, A., & Tosches, M. A. (2022). Cell-type profiling in salamanders identifies innovations in vertebrate forebrain evolution. Science, 377(6610), eabp9186. 10.1126/science.abp9186

Yoshino, J., & Tochinai, S. (2004). Successful reconstitution of the non-regenerating adult telencephalon by cell transplantation in Xenopus laevis. Development, Growth & Differentiation, 46(6), 523–534. 10.1111/j.1440-169x.2004.00767.x

Zwahlen, J., Gairin, E., Vianello, S., Mercader, M., Roux, N., & Laudet, V. (2024). The ecological function of thyroid hormones. Philosophical Transactions of the Royal Society B: Biological Sciences, 379(1898), 20220511. 10.1098/rstb.2022.0511

