## Supplementary Figures for "Glial cell states bias the regeneration of neuron types across the newt life cycle"

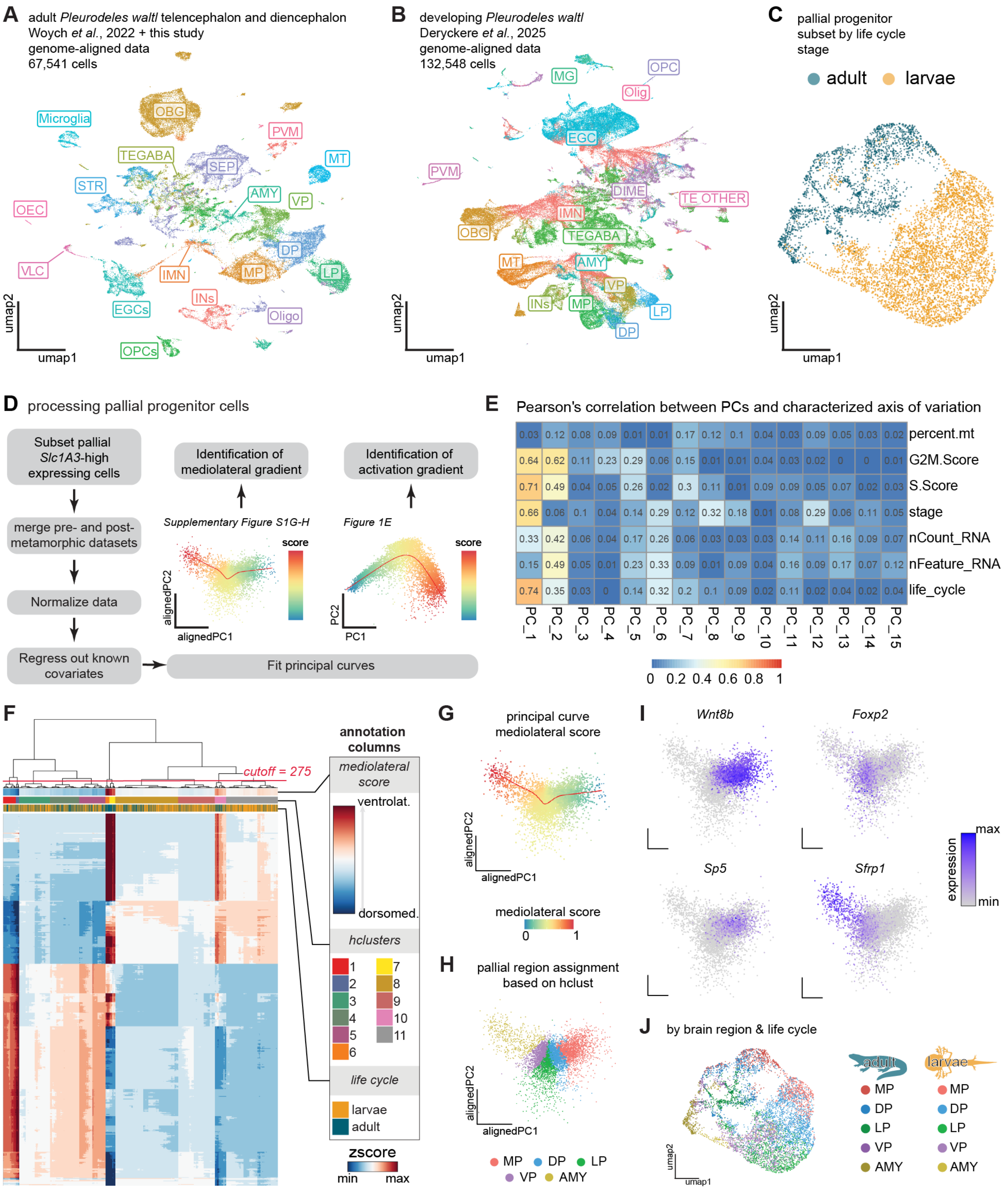


**Figure S1. Transcriptomic heterogeneity of *Pleurodeles* pallial progenitors.** (A) UMAP plot of the expanded scRNAseq dataset from the adult telencephalon (including data from Woych et al. 2022) aligned to the reference genome and annotated using label transfer (see Methods). (B) UMAP plot of the developmental scRNAseq dataset used in Deryckere et al., 2025 aligned to the reference genome and annotated by iterative label transfer (see Methods). (C) UMAP plot of the merged neural stem cell data (RGCs and EGCs, from larvae and adults) colored by life cycle stage. (D) Schematic representation of the analysis pipeline. (E) Heatmap showing Pearson's correlation between PCs and known biological and technical variables before regression. (F) Hierarchical clustering (Euclidean distance, ward.d2 method) and heatmap of the final progenitor dataset, showing 11 clusters at the cutoff (red line, h=275, see Methods). The heatmap shows genes with significant differential expression along the mediolateral axis principal curve, and clustering is based on the expression of the same genes. (G) alignedPCA reduction plot of the progenitors showing the cells colored according to the mediolateral score assigned with the principal curve. (H) alignedPCA reduction plot with cells colored by discrete pallial regions determined after clustering according to the mediolateral score. (I) alignedPCA reduction plots showing the expression of marker genes in the mediolateral axis. (J) UMAP plot of the merged larval and adult neural stem cells in C, colored by the pallial region of origin inferred from the mediolateral score analysis.

Abbreviations: AMY, amygdala; DP, dorsal pallium; EGCs, ependymoglia cells; IMN, immature neurons; INs, interneurons; LP, lateral pallium; MG, microglia, MP, medial pallium; MT, mitral and tufted; OBG, olfactory bulb granule cells; OEC, olfactory ensheating cells; Oligo, oligodendrocytes; OPCs, oligodendrocyte precursor cells; SEP, septum; STR, striatum; TEGABA, telencephalic GABAergic; TEOTHER, telencephalic other; VLC, vascular leptomeningeal cells; VP, ventral pallium.

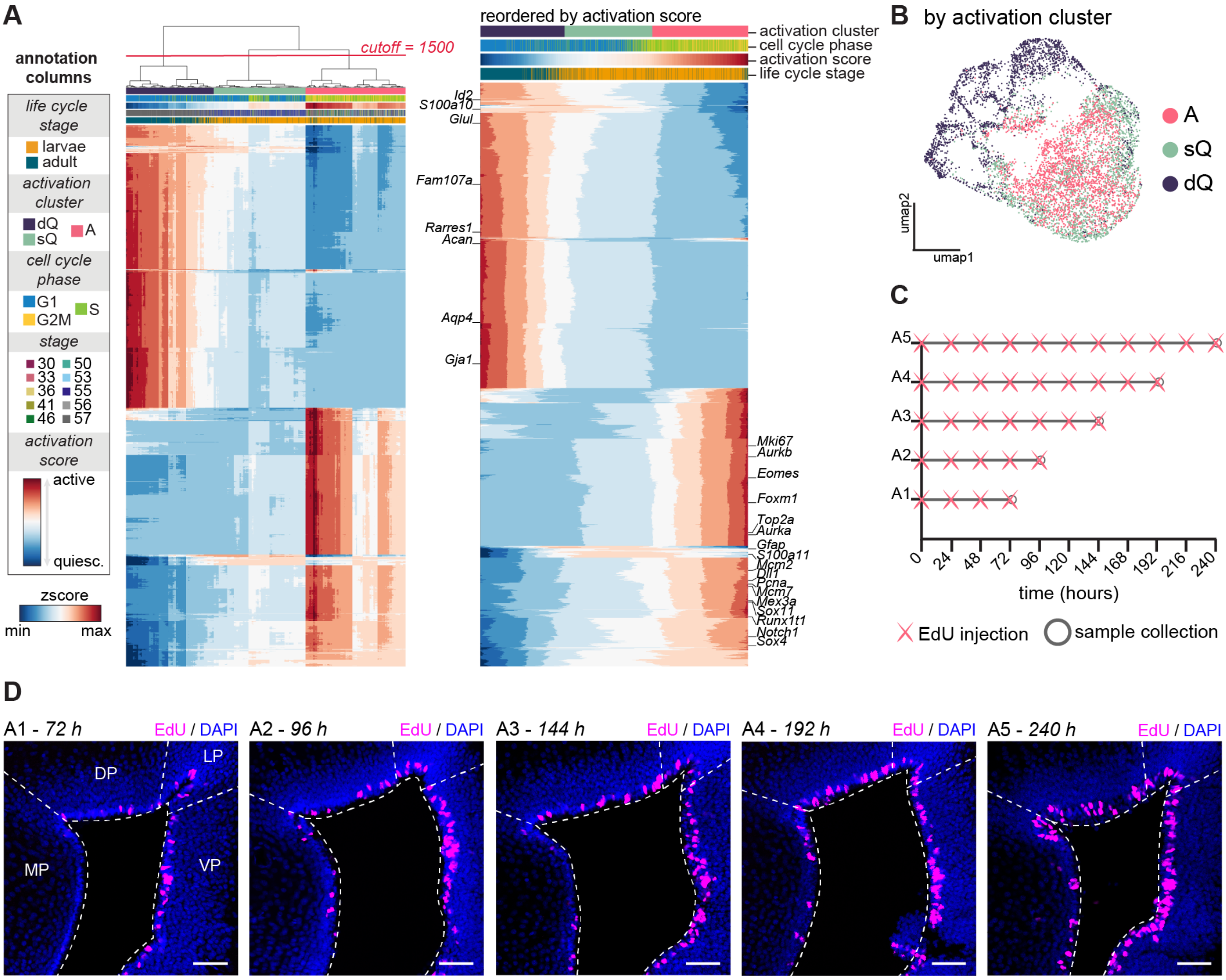


**Figure S2. Further analysis and validation of EGCs cell states across life cycle stages and pallial regions.** (A) Left: Heatmap and hierarchical clustering analysis (Euclidean distance, ward.d2 method) of the expression of 1971 genes with variable expression along the principal curve fitted in PC1/PC2, showing 3 clusters at the cutoff (red line, h=1500, see Methods). Right: Heatmap of the expression of the same 1971 varying genes in progenitor cells ordered by activation score from the principal curve. (B) UMAP plot of the merged larval and adult RGCs and EGCs, colored according to the activation clusters assigned with the activation score analysis. (C) Schematic representation of the cumulative EdU experiment. Adult animals matched by age and size received EdU injections every 24 hours, for a total of 4 (group A1) to 11 injections (group A5). Brains were collected two hours after the last EdU injection. (D) EdU labeling in the pallium of newts in the different experimental conditions described above (A1 to A5). Images show increasing incorporation of EdU in the EGCs of different pallial regions with increasing number of EdU injections, with a higher abundance of EdU+ cells in the VP and LP compared to MP and DP. Scale bar: 50 $\mu$m.

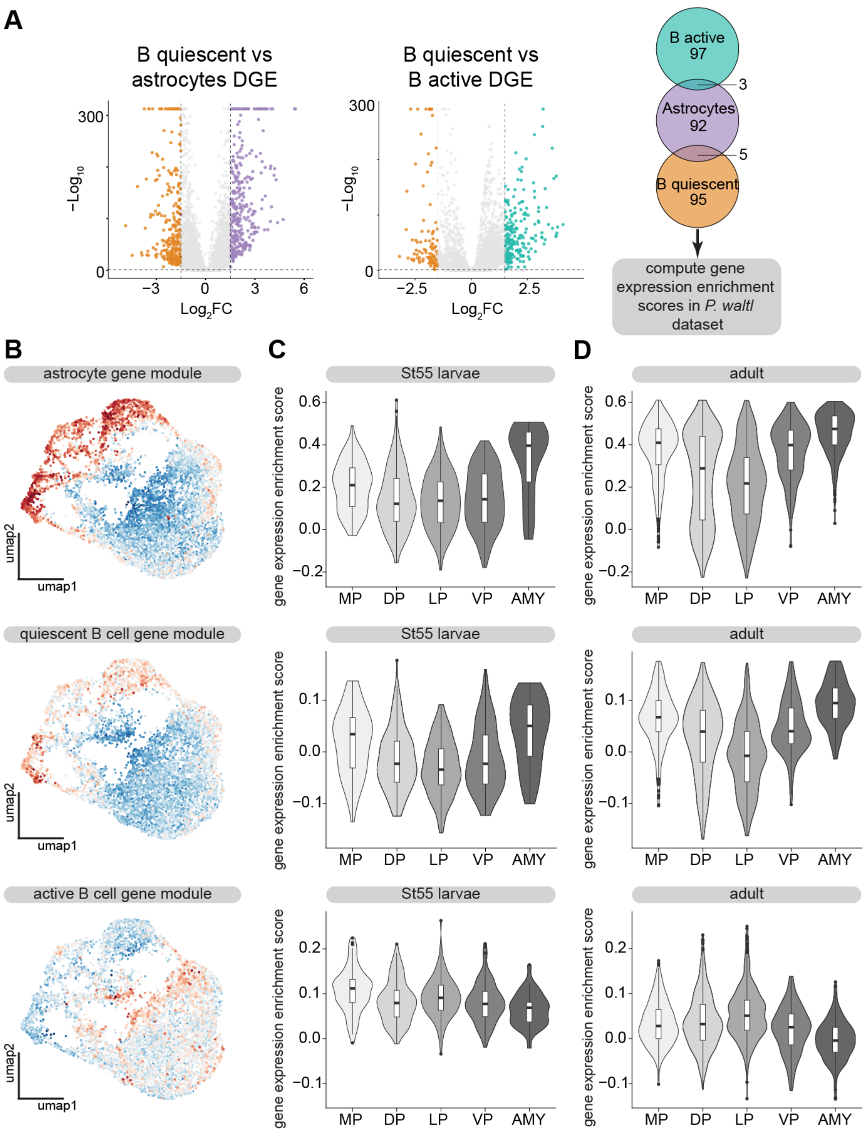


**Figure S3. Expression enrichment of mouse gene modules in the newt progenitor dataset.** (A) Volcano plots of the result of the differentially expressed genes (DEGs) computed from the mouse cell types: (Left) quiescent B cells *vs* astrocytes, (Middle) quiescent B cells *vs* active B cells from the Cebrian-Silla et al., 2025 dataset. Right: Intersection of the top 100 DEGs identified per comparison (see Methods), showing the total number of unique genes used in downstream analysis. (B) UMAP plots of the mouse gene module score enrichment in the newt dataset. (C) Violin plots for gene module score distribution in each pallial region in Stage 55 RGCs. (D) Violin plots for gene module score distribution in each pallial region in adult EGCs.

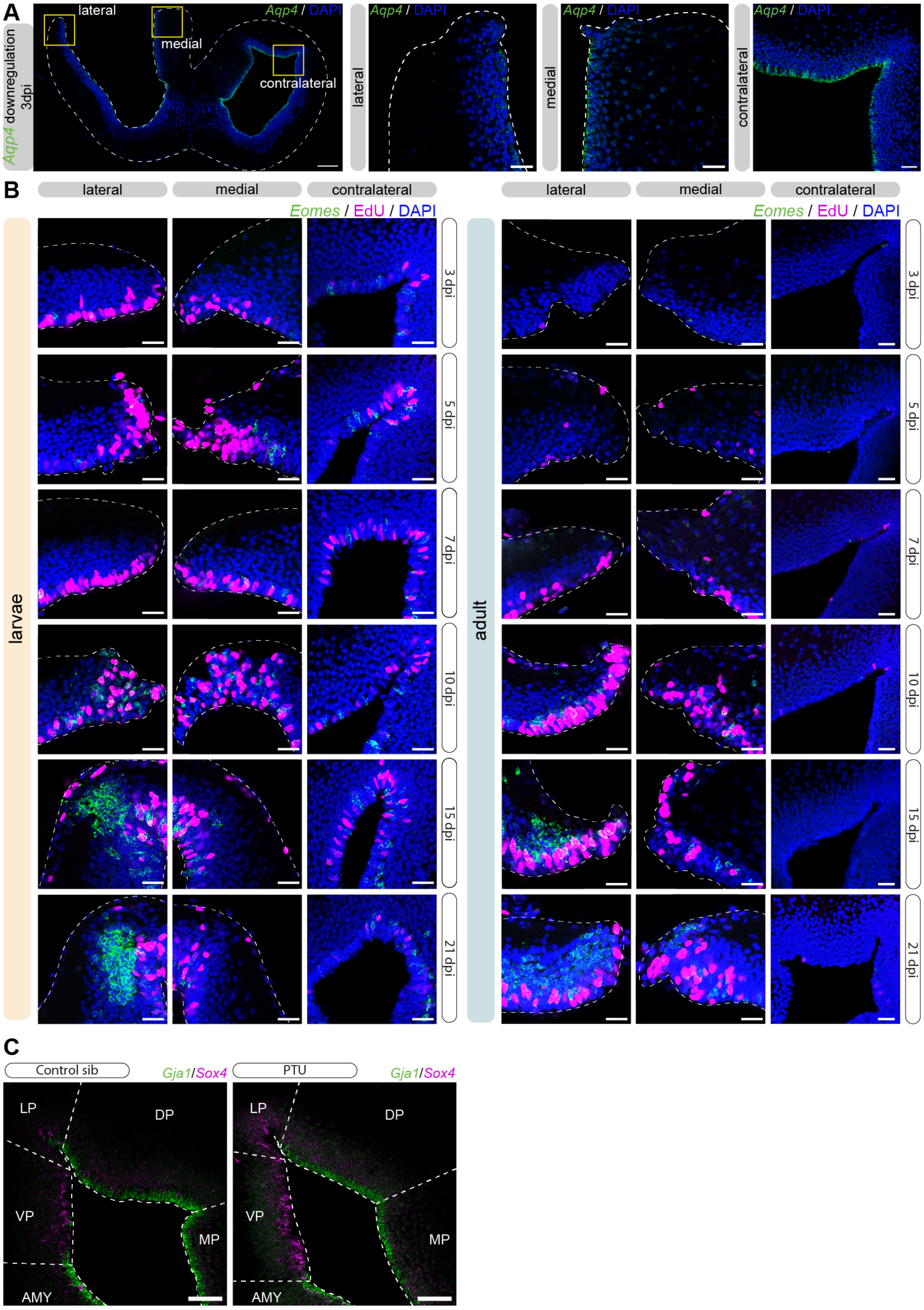
A
Aqp4 downregulation 3dpi
lateral
medial
contralateral
Aqp4 / DAPI
B
lateral
medial
contralateral
Eomes / EdU / DAPI
larvae
adult
3 dpi
5 dpi
7 dpi
10 dpi
15 dpi
21 dpi
C
Control sib
PTU
Gja1/Sox4
LP
DP
VP
MP
AMY

**Figure S4. Downregulation of quiescence, proliferation and IPCs in early stages after injury and PTU phenotype.** (A) Left: Overview of a mid-telencephalic coronal section of an adult brain at 3 dpi, stained with the deep quiescence marker *Aqp4*. Right: Magnifications of a coronal section showing the expression of *Aqp4* at 3 dpi at sites lateral, medial, and contralateral to the injury. Scale bar: 200 $\mu$m in the overview (left) and 50 $\mu$m for the magnifications. (B) Magnifications of coronal sections of the adult and larval pallium in the lateral and medial regions relative to the injury site and contralateral hemisphere at 3, 5, 7, 10, 15, and 21 dpi, showing proliferating cells (EdU+) and IPCs (*Eomes*). Scale bar: 50 $\mu$m. (C) Magnifications of coronal sections of the pallium in age-matched control animals and PTU-treated siblings showing deep quiescent EGCs (*Gja1*) and immature neurons (*Sox4*). Scale bar: 50 $\mu$m.

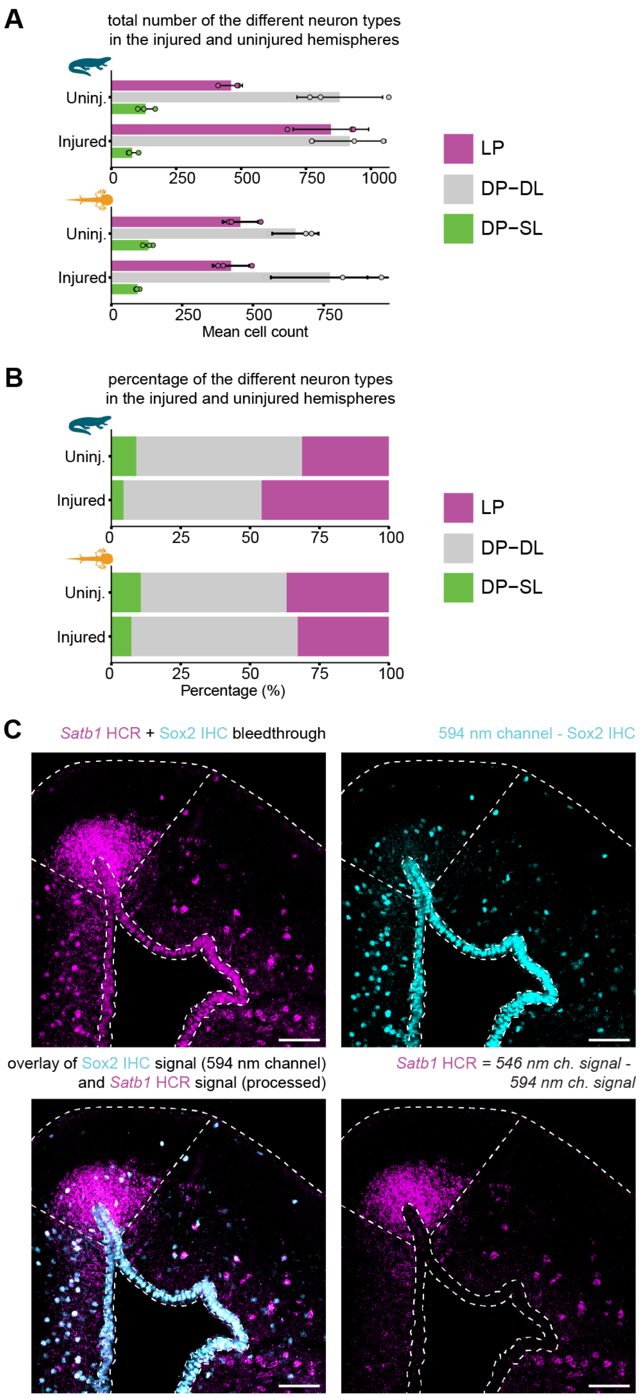


**Figure S5. Cell type quantification and image processing of regenerated pallium of larvae and adults.** (A) Barplot of total number of different neuron types (LP neurons, magenta; DP-DL, grey; DP-SL, green) present at 84 dpi in the injured vs. uninjured hemispheres of adults (top) and larvae (bottom). (B) Stacked barplot of percentage of different neuron types (LP neurons, magenta; DP-DL, grey; DP-SL, green) present at 84 dpi in the injured and uninjured hemispheres for adults (top) and larvae (bottom). Abbreviations: DP-DL, dorsal pallium deep-layer neurons; LP, lateral pallium; MP, medial pallium; VP, ventral pallium; DP-SL, dorsal pallium superficial-layer neurons. (C) Example of image processing to disambiguate *Satb1* HCR and Sox2 IHC signals. Top left: coronal section at mid-telencephalic level showing the 546 nm channel, corresponding to *Satb1* signal with bleedthrough Sox2 signal. Top right: the same coronal section showing the 594 nm channel, corresponding to the Sox2 signal. Bottom left: overlay of the Sox2 IHC signal (594 nm channel) and *Satb1* HCR signal (after subtracting the 594 nm channel from the 546 nm channel with Fiji). Bottom right: *Satb1* HCR signal after subtraction of the 594 nm channel.
